# C-Nucleosides Stabilize RNA by Reducing Nucleophilicity at 2’-OH

**DOI:** 10.1101/2025.07.23.666442

**Authors:** Dipanwita Banerjee, Lu Xiao, Pavitra S. Thacker, Jayanta Kundu, Muthiah Manoharan, Eric T. Kool

## Abstract

Nucleotides with carbon substitution for heteroatoms are common in biological and therapeutic RNAs. Important examples include the C-nucleosides pseudouridine and N1-methyl-pseudouridine; these modifications were reported to slow degradation of large RNAs, but the mechanism is unknown. We measured kinetics of spontaneous and enzymatic cleavage at a single bond of synthetically modified RNAs, and find that carbon substitution markedly reduces strand cleavage rates in RNA by both mechanisms. Studies of nucleophilic acylation reactions of RNAs and of small alcohols of varied p*K*_a_ suggest that reduced inductive effects resulting from carbon substitution for electronegative atoms results in both higher p*K*_a_ and lower nucleophilicity. The results provide insight into native transcriptome modifications as well as RNA therapies.

TOC Graphic

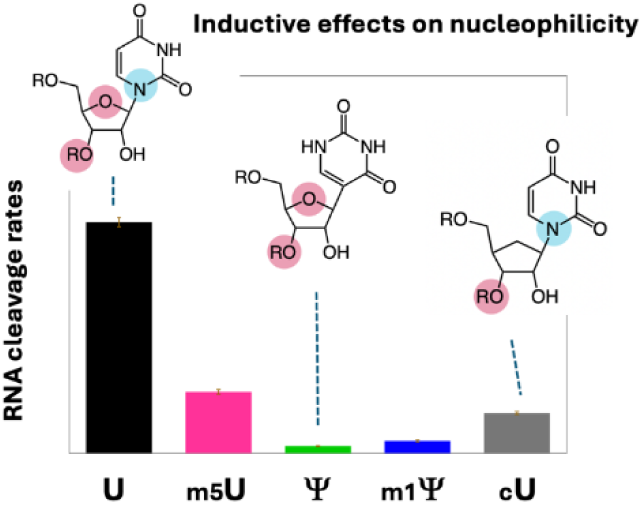

**Synopsis:** This study reveals mechanisms by which modifications found in native and therapeutic RNAs enhance RNA stability. Carbon substitution for nitrogen or oxygen results in reduced 2′-OH nucleophilicity.

## INTRODUCTION

Modified RNAs are under intensive development as drugs, genetic therapies, and vaccines. These bioactive species invariably incorporate numerous nucleobase and sugar modifications to provide enhanced stability to degradation and reduced innate immune responses.^1–3^ Many of these modifications are inspired and adopted from modifications first identified in naturally occurring RNAs.^4–6^ Over 150 chemical modifications have been characterized in native RNAs, adding a regulatory layer to RNA biology by affecting cellular processes such as gene expression,^4^ splicing,^5^ and translation.^6,7^

One of the chief motivations for incorporating ribonucleotide modifications in RNA therapies is the fact that the biopolymer is highly susceptible to degradation, resulting in strand cleavage due to spontaneous and enzymatic processes. This strand cleavage directly involves the 2’-hydroxyl (2’-OH) group of the ribose sugar, which undergoes intramolecular nucleophilic attack at the neighboring phosphate, breaking the 3’-5’ phosphodiester bond. The non-enzymatic intramolecular transesterification of RNA is known as “in-line” cleavage (Figure 1).^8,9^ This nucleophilic attack occurs spontaneously under nonenzymatic conditions, especially at elevated pH, where deprotonation of the 2’-hydroxyl group generates a highly reactive 2’-oxyanion (Figure 1C),^10^ or is facilitated by nuclease enzymes that provide general acid/base catalysis and promote the required conformation changes for this attack.^11,12^ Chemical modification or replacement of the 2’-OH can be beneficial, extending the lifetime of bioactive RNAs in the cell.^13,14^ Studies of modified nucleotides have provided important outcomes in improving RNA therapies and vaccines for increasing target specificity and reducing immunogenicity.^2^ However, mechanistic understanding of the effects of nucleotide modifications on RNA degradation is less well explored. Such understanding can provide both basic insights into existing naturally occurring modifications in RNA, as well as inspire improved designs for future RNA therapies.

**Figure 1.**
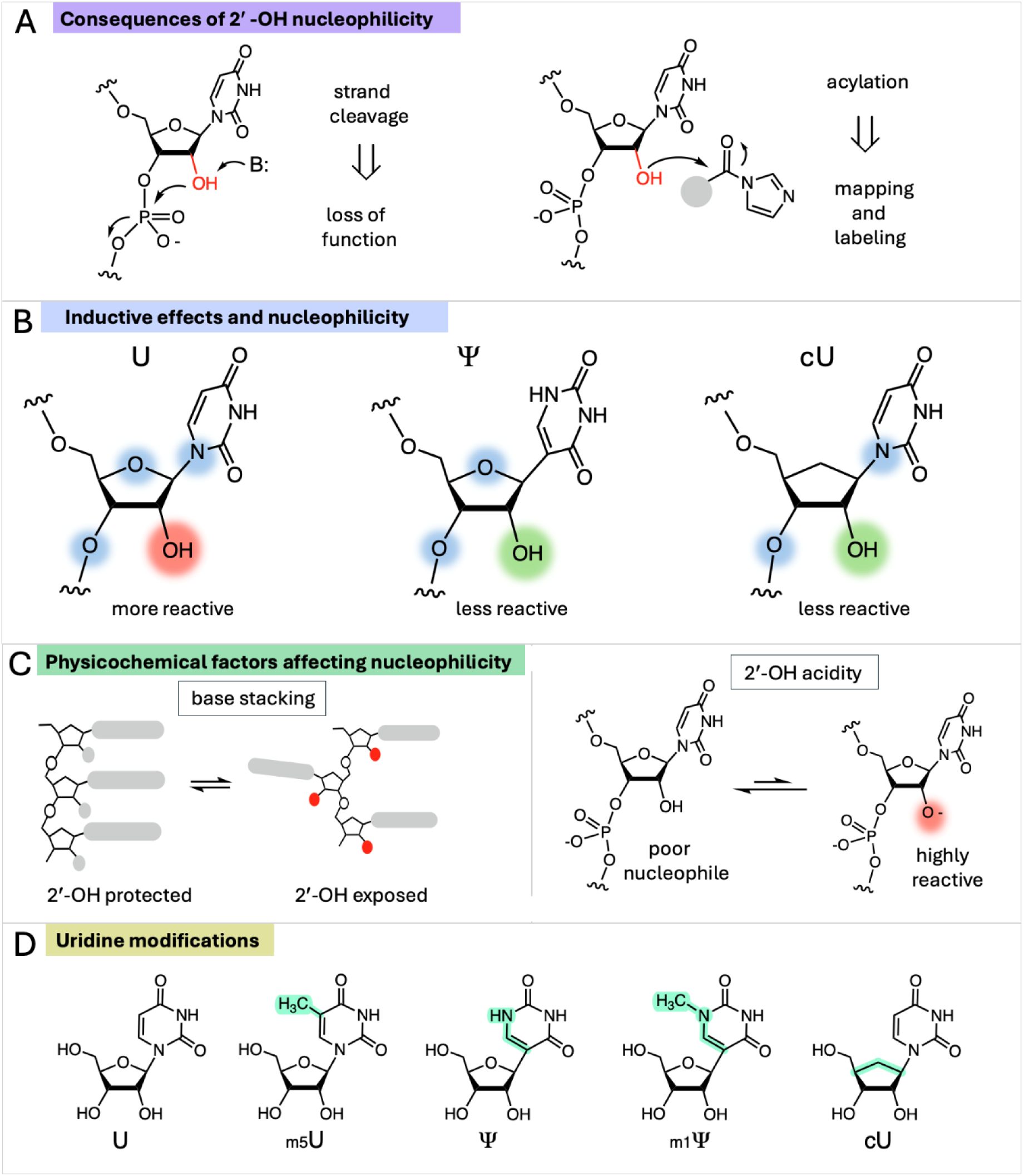
The effects of pyrimidine nucleotide modifications on RNA stability. (A) Modifications that alter the nucleophilic reactivity of the 2′-OH group can affect RNA strand cleavage rates and acylation rates. (B) We hypothesize that modifications that reduce the number of electronegative N/O atoms (highlighted in blue) within two bonds of 2′-OH lower its acidity (increase its p*K*_a_), as is known for small-molecule alcohols. This inductive electron-withdrawing effect of N/O stabilizes the oxyanion form of the hydroxyl; conversely, carbon substitution for these electronegative atoms decreases the population of the anionic form. (C) Strongly stacking uridine analogs test the effects of base stacking on 2′-OH reactivity; increased stacking stabilizes the helical conformation, which has lower nucleophilic reactivity. Analogs with altered acidity will influence the fraction of the highly reactive oxyanion form. (D) Structures of uridine and analogs employed in this study, with differences highlighted in green.

Modified nucleotides can potentially protect RNA from degradation by multiple mechanisms. One clear mechanism is direct modification or replacement of the 2’-OH group to provide a chemical block to intramolecular nucleophilic attack. Examples of this strategy include 2’-O-methyl groups, 2’-O-acyl groups, and 2’-fluoro groups, all of which completely prevent cleavage. However, other nucleotide modifications more remote from the 2’-OH group (Figure 1) also have the potential to enhance stability against strand cleavage. Indeed, non-natural base modifications have begun to be employed recently to explore how nucleobases may interact with phosphodiester linkages to alter phosphodiester cleavage.^15,16^ Most relevant to the current work, Kariko *et al*. reported that replacement of uridine by pseudouridine (Ψ) in mRNAs provides a measure of stabilization against degradation in cells; however, the mechanism by which this occurred was not determined.^17^ This modified nucleoside is an isomer of uridine in which the base is attached to ribose through carbon rather than nitrogen, resulting in a C-nucleoside (Figure 1B). A more recent study showed that Ψ replacement of U enhances stability of an mRNA to spontaneous degradation.^18^ In general, studies of the degradation of unmodified RNAs have shown that single-stranded, flexible regions are hotspots for cleavage.^18,19^ Thus it is plausible that base modifications that thermodynamically enhance stacking and helicity have the potential to indirectly stabilize RNAs against strand cleavage. Interestingly, thermodynamics studies have shown that Ψ stabilizes RNA helices moderately relative to uridine,^20^ providing one possible mechanism for suppressing strand cleavage.

Here we also consider a different and previously unrecognized mechanism for RNA strand stabilization by carbon-substituted nucleotides, involving differences in the inherent nucleophilicity of the 2’-OH group. Studies of the nucleophilicity of the 2’-OH group toward acylating agents at physiological pH suggest that the main reactive form of this hydroxyl is the oxyanion (Figure 1C),^21^ which is rare at pH 7.4 but rapidly equilibrates. The p*K*_a_ of the 2’-OH group in mononucleotides is approximately 12.5,^22^ *ca*. 3 p*K* units lower that of than reference alcohols such as ethanol or isopropanol, and measurements of pH-dependent cleavage rates in RNA models have suggested a p*K*_a_ of *ca.* 13.6.^23^ We have suggested that the relative acidity of the 2’-hydroxyl group in canonical RNA is at least in part due to inductive effects of three highly electronegative atoms in proximity: the 3’-oxygen, the 4’-oxygen, and the N1 nitrogen of the base (Figure 1B).^24^ In this light, we note that C-nucleosides commonly employed in therapeutic RNAs (Ψ and N1-methyl Ψ (m1Ψ)) have one of these three electronegative atoms replaced by carbon. Thus, it seems possible that the nucleophilic reactivity of the 2’-OH group might be reduced in such C-nucleosides as a result of lower acidity; this might be expected to reduce rates of reactivity with electrophiles including both external acylating agents as well as the neighboring phosphodiester group (Figures 1B,C). To date, no kinetics data for these nucleosides have been available for either of these forms of reactivity.

In sum, multiple physical and chemical factors of nucleotide modifications have the potential to affect RNA strand stability against spontaneous and enzymatic cleavage. In this study, we analyze these factors in short RNAs by use of specifically placed carbon modifications (Figure 1D). We compare reactivity of C-5 methylated nucleotides, as such methylation is documented to enhance stacking,^25^ to unmethylated analogs, and to test the effects of the absence or presence of electronegative atoms on nucleophilicity and cleavage, we study RNAs with C-nucleosides Ψ and m1Ψ, as well as cU, a carbacyclic uridine analog.^26^ Nucleophilic reactivity is assessed with spontaneous and enzymatic bond cleavage measurements as well as with acylation reactions.

Overall, our results show that methyl modifications that alter base stacking have mixed and moderate effects on nucleophilicity and strand cleavage. However, we find that the 2’-OH group of the canonical N-nucleoside uridine is significantly more reactive as a nucleophile than the analogous hydroxyl of C-nucleosides, resulting in considerably higher rates of acylation and of nonenzymatic phosphodiester bond cleavage. Enzymatic cleavage by ribonuclease A and other nucleases is also strongly suppressed at Ψ and m1Ψ as compared with U. Studies of the reactivity of small alcohols of varied p*K*_a_ suggest that a major contributor to this difference in nucleophilicity is inductive effects on 2’-OH acidity as a result of C versus N substitution.

Taken together, our studies provide new mechanistic insight into the effects of native RNA modifications on their biological properties, and provide information that may aid in future improvements in the stability of therapeutic RNAs.

## RESULTS AND DISCUSSION

### Shielding of RNA from spontaneous degradation by nucleotide modifications

While our primary studies were carried out with short synthetic oligoribonucleotides, we first confirmed that a pseudonucleotide replacement for uridine can stabilize mRNA to spontaneous degradation under our assay conditions (Figure 2A). A previous study reported this for an mRNA;^17^ here we sought to confirm the effects of m1Ψ under our conditions using the enhanced green fluorescence protein (eGFP) mRNA context. Although this replacement occurs at less than 20% of the nucleotides of this RNA, our results demonstrate a ∼2-fold increase in the half-life of the full-length RNA, indicating significant stabilization (Figure 2A). This is consistent with the previous measurements of mRNAs containing Ψ and m1Ψ,^18^ confirming that they provide a degree of stabilization in a large RNA even as minority substitutions among canonical nucleotides.

**Figure 2.**
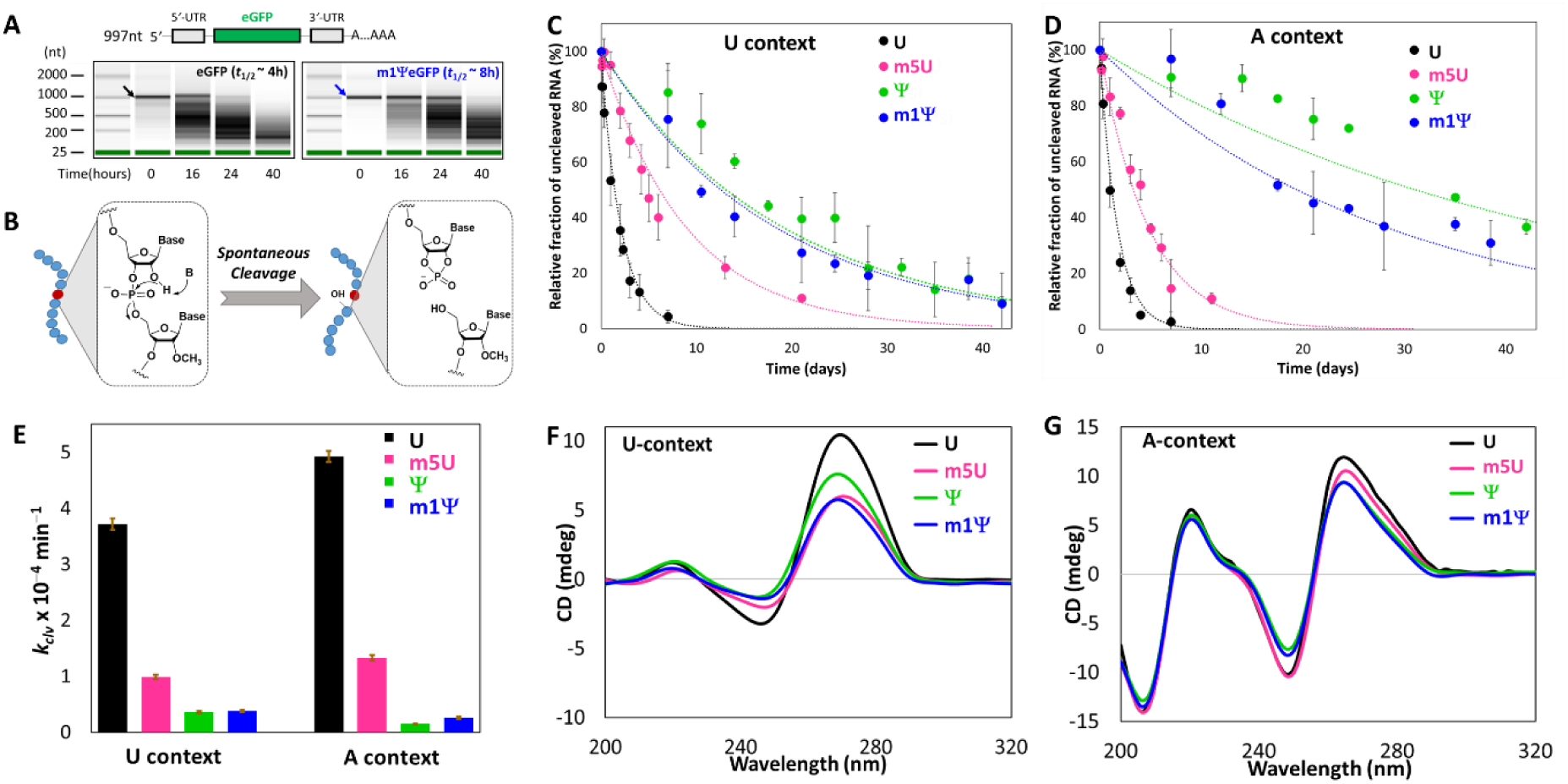
Base modifications shield RNA from spontaneous degradation. (A) mRNA stabilities assessed over time (water, 37 °C) for unmodified eGFP RNA and m1Ψ-modified eGFP RNA measured by capillary electrophoresis. (B) Molecular mechanism for in-line cleavage of scissile phosphodiester bond in RNA oligonucleotides. Nucleotides with unstable and stable 3′-phosphodiester bonds are colored red and blue, respectively. (C, D) Plots of the cleavage of 11 nt RNAs after the central U or modified nucleotide by spontaneous cleavage at 37 °C in CHES buffer solution (pH 10.2) in U and A contexts, respectively. Data are averages from 3 replicates with error bars showing standard deviations. (E) Comparison of the rate constants (k_clv_) derived from the spontaneous degradation shown in C and D. (F, G) Circular dichroism (CD) spectra for 11 nt RNAs with U and A contexts, respectively, with a central U or analog.

For RNA polymers, all internucleotide linkages can simultaneously undergo cleavage, making quantitative measurements of rates challenging. To determine the effects of nucleotide modification on individual phosphodiester bond cleavage, we designed short (11 nt) RNA oligonucleotides (Table S1) containing only one scissile phosphodiester bond at the 3*’* side of uridine and analogs (Figure 2B), with the adjacent nucleotides 2*’*-*O*-methylated to block cleavage at all other positions. Importantly, 2*’*-*O*-methylated RNAs retain a helical conformation nearly identical to that of unmethylated RNA oligonucleotides.^27^ To investigate the effect of base stacking context for the central uridine analogues, we studied contexts of polyU and polyA sequence (Table S1). Adenine is in some contexts the most strongly stacking base in RNA, while uracil is the least.^28^ Guanine also stacks strongly but introduces additional complexity due to Hoogsteen pairing, resulting higher order non-canonical structure formation. UV-monitored thermal melting studies in buffer (pH 7.0) confirmed that no significant higher-order intramolecular or intermolecular structure was formed by the selected RNA oligomers (Figure S1).

Kinetics of single bond cleavage events were assessed from base-catalyzed cleavage reactions of the oligonucleotides, monitored with gel electrophoresis. The mechanistic pathway of cleavage of RNA polymers involves in-line phosphoester transfer (Figure 2B).^8,29^ The RNAs were incubated at 37 °C in a solution of 50 mM CHES (pH 10.2) containing 10 mM MgCl_2_, similar to prior studies;^23^ A Mg²⁺ concentration of 10 mM was essential to achieve efficient RNA cleavage.^23^ After quenching the reactions, the fraction of strand cleavage was quantified by denaturing polyacrylamide gel electrophoresis (PAGE) (Figure S2). Plots of intact RNA fraction *vs* time were fit with first-order exponential decay curves (Figures 2C,D and S3).

Strand cleavage rate constants (*k*_clv_) are given in Table 1, and are compared graphically in Figure 2E. Data were obtained for uridine and four uridine analogs in two contexts. Comparing the adjacent sequence contexts of polyU versus polyA, we find moderate and mixed effects; for the N-nucleotides U and m5U, the rate constant for cleavage was 1.3-fold lower in the U context relative to the A context. However, for the C-nucleotides Ψ and m1Ψ, the reverse was true, with slightly slower cleavage occurring in the A context. Previous studies have shown that sequence context can affect RNA cleavage rates significantly.^30^

**Table 1.**
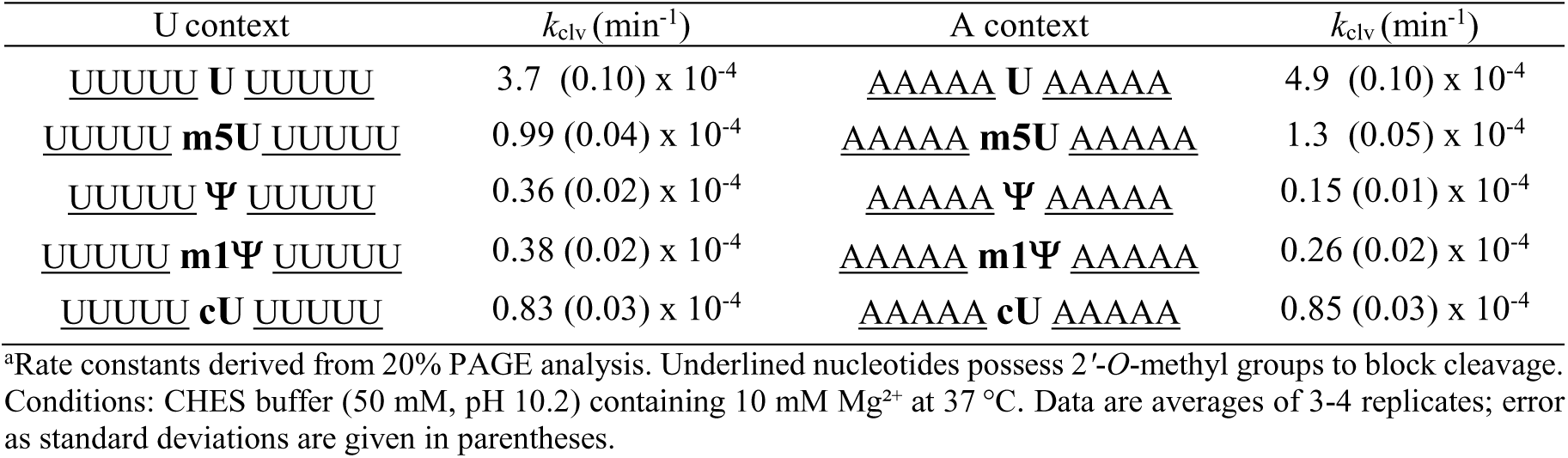
Rate constants for spontaneous cleavage at 3′ of uridine and analogues in oligoribonucleotides.^a.^

Adjacent (nearest-neighbor) bases affect local RNA structure and stability by differential stacking free energies.^20^ Methylation of uridine (in m5U) enhances base stacking and stabilizes helical conformations of RNA,^25^ and *N*1-methylation of Ψ similarly stabilizes base stacking^.6,31^ m5U and m1Ψ are both known in therapeutic RNAs, as is the structurally related methylation of cytosine in m5C. Here we find that, as was seen for the varied adjacent base context, methylation also has moderate and conflicting effects. C-5 methylation of uracil results in nearly identical 3.6-3.8-fold greater stabilization of the internucleotide linkage against cleavage in both sequence contexts. In contrast, analogous methylation of Ψ results in either no effect or a small enhancement of cleavage rate.

Importantly, the data also allow a comparison of the effects of C-nucleoside to N-nucleoside substitution on the cleavage rates of the short RNAs. Results were consistent across context and methylation status, and were considerably greater in magnitude; all C-nucleoside substitutions result in strong stabilization against strand cleavage. Substituting Ψ for U substantially stabilizes the RNA phosphodiester linkage against cleavage, resulting in 10-fold (U context) to 32-fold (A context) reductions in kinetic rate. Similarly, substitution of m1Ψ for m5U results in 2.7-fold to 5.3-fold drops in strand cleavage rates. Thus, the C-nucleoside substitution results in strong suppression of in-line attack on the phosphodiester by the 2*’*-OH group. While a previous study showed stabilization by Ψ of partially-substituted RNAs from spontaneous degradation,^18^ the current data provide quantitative measures of the effect on a specific bond.

Structural studies of the nucleosides indicate that the pseudouridine and N1-methylpseudouridine modifications alter sugar conformation slightly; Ψ and m1Ψ have 48:52 and 46:54 North/South conformational preferences, respectively, shifted by a small degree from that of uridine (53:47 North/South).^32^ The Ψ modification also moderately enhances base stacking relative to U depending on sequence context.^20,33^ CD spectra of the oligonucleotides (Figures 2F,G) showed evidence for moderate reduction of helicity in the U context by the base modifications. However, no changes in helicity were seen for the A context. Our data show that C-nucleoside substitution has a stronger suppressive effect on strand cleavage in the A context, where no change in overall helicity is seen, suggesting that global changes in oligomer conformation are unlikely to be the source of the protective effect. Thus, the data suggest a more localized explanation for most of the lower reactivity of 2*’*-OH groups in Ψ and m1Ψ.

### Base modifications protect RNA from enzymatic degradation

To test whether enzymatic bond cleavage is also affected by uridine nucleotide modifications, we carried out quantitative measurements of cleavage with ribonuclease A, one of the most well-studied nuclease enzymes.^34^ RNase A is a metal ion-independent pancreatic ribonuclease,^35^ and its catalytic activity involves two histidine residues for general base/acid catalysis (Figure 3A).^36^ Although RNase A-catalyzed degradation follows a similar mechanistic pathway as in-line uncatalyzed cleavage, it provides ∼10^11^-fold rate acceleration.^37^ RNase A prefers to cleave the phosphodiester bond to the 3*’* side of pyrimidine nucleotides and 5*’* to purines,^34^ enabling our 11 nt oligomers in the polyA context to act as appropriate enzyme substrates. We are aware of no prior studies testing the effects of C-nucleoside structure or methylation on RNase A nuclease activity.

**Figure 3.**
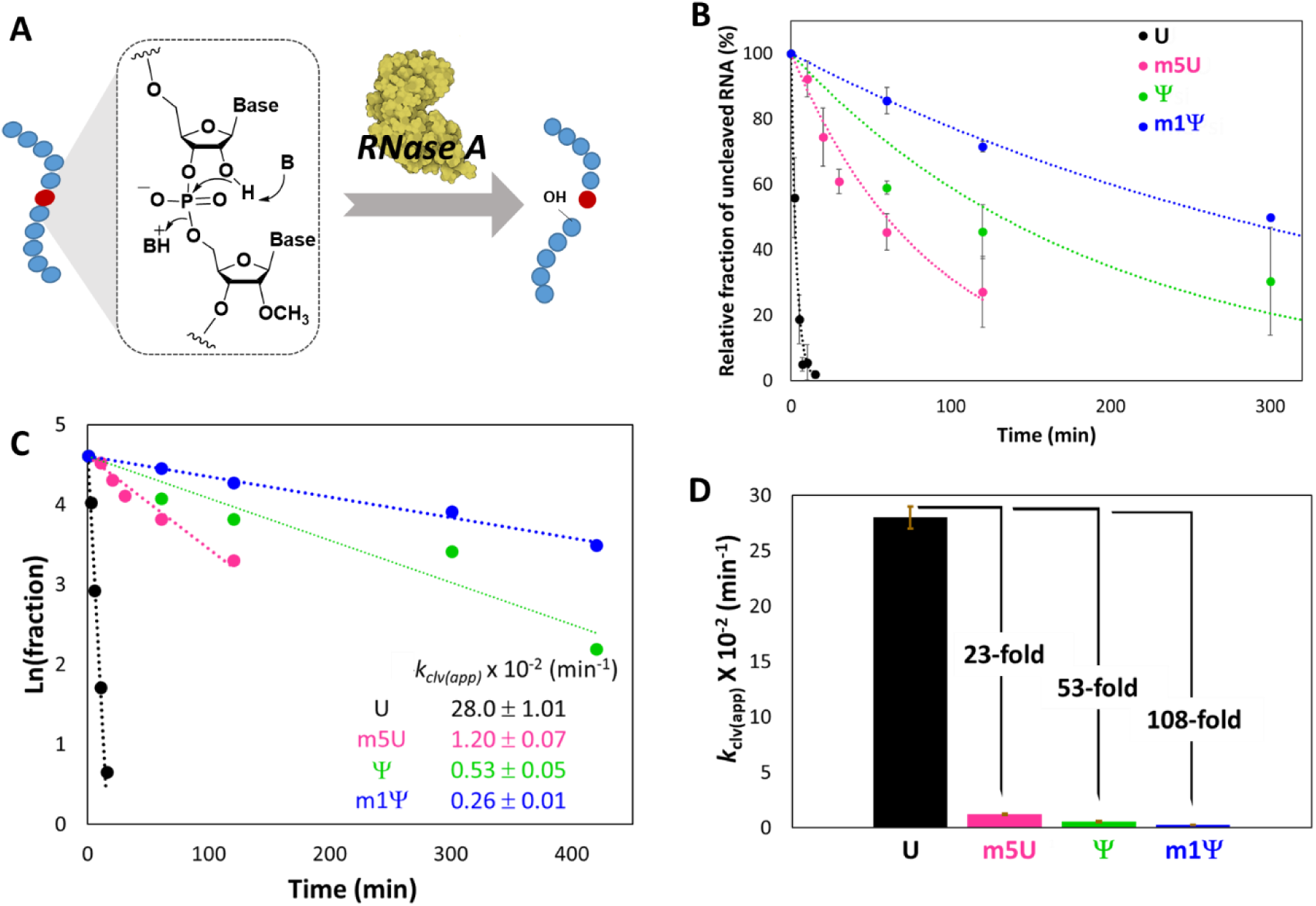
Modifications protect RNA from ribonuclease cleavage. (A) Scheme of substrates employed in RNase A catalyzed cleavage of a specific internucleotide linkage in RNA oligonucleotides. (B) Plots showing decay of RNA phosphodiester by enzymatic cleavage at 37 °C with RNase A for oligonucleotides containing U, m5U, Ψ, and m1Ψ with time. (C) Linear plots of Ln(remaining RNA fraction) vs time for unmodified and modified uridine-containing RNAs. Negative slopes indicate pseudo-first-order rate constant, from which second-order rate constant was obtained. (D) Comparison of apparent 2^nd^-order rate constant values of RNase A-catalyzed degradation (*k*_clv(app)_) for A-context oligonucleotides (see Table 1). Data were obtained from 3 replicates with error bars showing standard deviations.

We measured bond cleavage for the 11 nt sequences in the polyA context with comparison of U, m5U, Ψ, and m1Ψ. Due to very high enzymatic activity (Figure S4A), we used dilute RNase A solutions (10 ng mL^-1^ or 0.73 nM) with a large excess of RNA substrate to follow initial rates of cleavage at 37 °C in 50 mM Tris-HCl buffer (pH 7.4). The relative fractions of uncleaved oligonucleotide were quantified with denaturing PAGE (Figure S4) and plotted against time (Figure 3B). The *K*_M_ of RNase A has been reported as 1100 μM;^38^ as our concentrations were far below this, the data were fit well with pseudo-first-order kinetics (Figures 3B,C). Apparent second-order rate constants of RNase cleavage (*k*_clv(app)_) were derived from the linear pseudo-first-order plots (Figure 3C), and data are plotted in Figure 3D. The results show that both base methylation and C1-substitution decrease ribonuclease cleavage rates markedly. Methylation of U was associated with a 23-fold slower cleavage rate, but a 2.1-fold faster cleavage rate for m1Ψ compared to at Ψ. Most notably, C-nucleoside substitution had an even larger effect on cleavage, slowing strand scission by factors of 52 fold in the unmethylated case and 4.7 fold in the methylated case. Thus, the data show that both structural carbon substitutions have a strongly suppressing effect on a human ribonuclease-mediated RNA degradation.

Structures of RNase A with nucleotide substrates show close contacts of the enzyme with the pyrimidine base, explaining its preference for cleavage after a pyrimidine nucleotide in RNA.^39^ It is possible that methylation sterically inhibits interactions with the preferred substrate, although more detailed kinetics studies will be needed to shed light on this possibility. Regardless of mechanism, the results suggest that naturally occurring C-5 methylation of RNA pyrimidines may have a similar suppressive effect on cleavage of the 3*’*-phosphodiester bond. Although 5-methylC is known in mRNAs, and has been shown to prolong cellular lifetimes of the RNA,^40^ its effect on a specific nuclease enzyme has yet to be studied.

To evaluate the broader impact of C-nucleoside substitution on ribonuclease-mediated cleavage, we examined the activities of endonucleases RNase 4 and RNase 1, which are abundantly expressed in the human pancreas and serum, respectively. RNase 4 preferentially cleaves 3′ to U, while RNase 1 exhibits broad specificity for all nucleobases. In both cases, replacement of U by the C-nucleoside pseudouridine (Ψ) led to a pronounced suppression of enzymatic cleavage (Figure S5), with 110-fold and 13-fold reductions in *k*_clv(app)_ for RNase 4 and RNase 1, respectively. Notably, the extent of cleavage inhibition by Ψ varied significantly across the ribonucleases, potentially reflecting different degrees of hydroxyl deprotonation at the rate-limiting step, or distinct nucleobase binding interactions. Collectively, these findings reveal that Ψ broadly suppresses ribonuclease-catalyzed strand cleavage, offering mechanistic insights into its stabilizing role in RNA. Given the frequent occurrence of Ψ in natural RNAs, it is clear that one biological effect is likely to be suppression of enzymatic degradation rates at the bond 3*’* to the nucleotide. For RNase A, it seems difficult to attribute structural effects in the enzyme/RNA complex to explain this relatively large effect, since Ψ is no larger than U and adopts a very similar conformation. Thus, we hypothesize a local chemical effect of pseudouridine C-nucleoside substitution on reducing the inherent nucleophilicity of the attacking 2*’*-OH group (see below).

### Carbon substitution in the ribose sugar also slows strand cleavage

The above data show that carbon substitution for electronegative N at the C1*’* carbon of ribose can strongly reduce strand cleavage by spontaneous or enzymatic processes. We hypothesize that this replacement lowers nucleophilic reactivity of the nearby 2*’*-OH group by reducing the inductive effect of nearby electronegative atoms that stabilize the anionic form of the hydroxyl (Figures 1B,C). Although this was tested above with two C-nucleosides, we sought a further test of this hypothesis by replacing one of the other electronegative atoms in the vicinity of 2*’*-OH. To this end, we employed the recently described carbacyclic uridine analog cU (Figure 1),^26^ which has a chemical structure identical to U but replaces the C4*’* ring oxygen with carbon. Importantly, this substitution resides the same number of chemical bonds from the 2*’*-OH as the carbon substitution in the pseudonucleosides (Figure 1), which is relevant because inductive effects depend strongly on the number of intervening chemical bonds.^41^ We prepared oligoribonucleotides having the same polyU and polyA contexts as the other modifications, and again measured rates of spontaneous cleavage (Figure 4). Comparison of strand scission rates again revealed a clear shielding effect of the substitution on the strand cleavage with cU. Carbacyclic uridine resulted in 4.5-fold and 5.8-fold decreases in *k*_clv_ compared to unmodified uridine in the pyrimidine and purine contexts. This effect is similar in magnitude to the suppressive effect of C-for-N replacement in the pseudonucleotides (Figure 2E). CD spectra revealed that the RNAs containing cU exhibit moderately or slightly reduced global helicity as the analogous RNAs with U, respectively (Figure S6), suggesting that the stabilization in this case was not due to a large change in conformation, but is instead localized near the modification.

**Figure 4.**
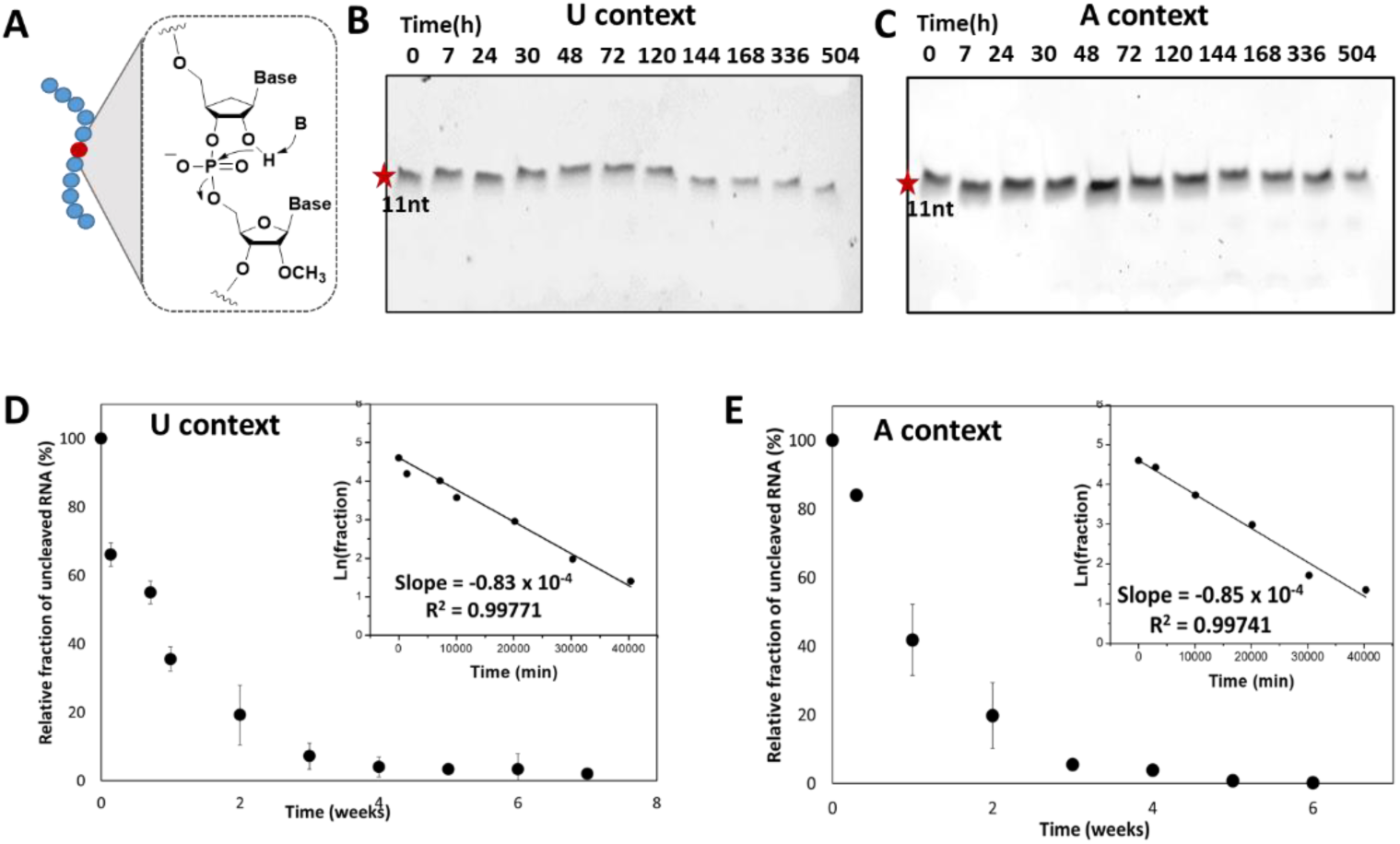
Shielding of RNA from spontaneous degradation by carbacyclic uridine modifications. (A) Molecular mechanism for in-line cleavage of scissile phosphodiester bond in carbacyclic uridine modified 11 nt RNA oligonucleotides. Nucleotide with cleavable 3*’*-phosphodiester bond is colored red. (B,C) Representative images of 20% denaturing PAGE analysis for spontaneous cleavage of cU RNAs in U and A contexts, respectively, after incubating substrate oligonucleotides in CHES buffer (pH 10.2) solution with Mg^2+^ at 37 °C. (D, E) Decay curves from band intensities; inset plots depict the first-order decay behavior, with slopes yielding rate constants *k*_clv_ (min^-1^). Data were obtained from 3 replicates with error bars showing standard deviations.

### Carbon substitutions reduce nucleophilicity as measured by 2*’*-OH acylation

The above data show that carbon substitutions near the 2*’*-OH group of a ribonucleotide in RNA lower the nucleophilicity of this hydroxyl for the neighboring phosphate group. To evaluate nucleophilicity of this group in a different context and at neutral pH, we employed a different electrophile to test. The 2*’*-OH group of RNA can be modified by acylating agents in water, a fact that has been widely used in mapping RNA structure in SHAPE methods^42^ as well as in strategies for high-yield labelling and stabilization of RNA.^43^ Thus, we used an acylation reaction here to further evaluate the nucleophilicity of 2*’*-OH (Figure 5A). Notably, no prior study has to our knowledge quantified absolute acylation kinetics in full RNA strands, the results of which can be relevant both to RNA structure mapping and to development of new methods and reagents for RNA labelling.

**Figure 5.**
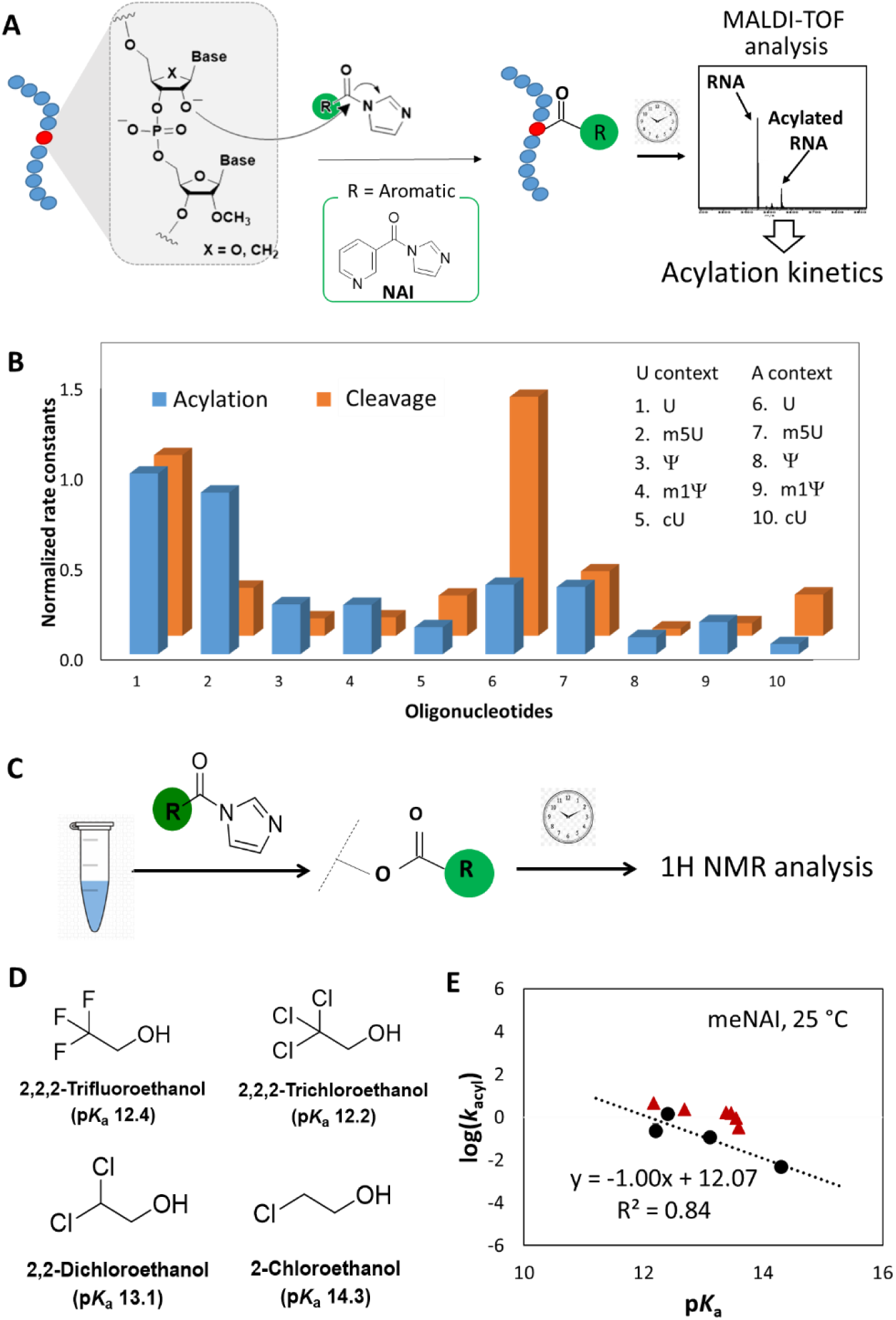
Effect of 2*’*-OH nucleophilicities on acylation. (A) Molecular mechanism and experimental workflow of RNA 2′-OH acylation with 10 different unmodified and modified oligonucleotides (Table S1). MALDI-TOF quantifies the rate constants of the acylation using the initial rates method. (B) Comparison of normalized rate constants for the acylation of modified nucleotides in the RNA oligonucleotides with corresponding cleavage rate constants. Normalization was done with respect to the value obtained for unmodified U in the U context. (C) Experimental workflow to study kinetics of acylation for alcohols with NAI at 37 °C or meNAI at 37 °C (pH 7.4). (D) Molecular structures of alcohols with p*K*_a_ range (12-15) at 25 °C to study the correlation of nucleophilicity with p*K*_a_. (E) Plot of the logarithms of rate constants for primary alcohols (black circle) against reported p*K*_a_, bracketing likely RNA p*K*_a_ values. Previously reported log(*k*_Acyl_) and p*K*_a_ values for nucleotides are also shown (red triangles).

As described above, these RNA sequences (Table S1) contain only one free 2*’*-OH available for acylation, simplifying the measurement of acylation yields and rates. Reactions were quantified from relative peak intensities measured by MALDI-TOF mass spectrometry over time (Figure 5A). Rate constants of acylation reaction (*k*_acyl_) were determined with an established RNA acylimidazole reagent (NAI), employed previously to study kinetics of acylation with mono- and di-nucleotides.^21^ Data are shown in Table 2 and Figure S7. Comparison of rates (Figure 5B) clearly distinguishes the effects of carbon substitution and base stacking. Substitution of U with Ψ and m5U with m1Ψ led to 3.6-fold and 3.2-fold reductions in *k*_acyl_, respectively, whereas no comparable change was observed due to the methylation of U and Ψ in the polyU context. This suggests that there is a negligible effect of base stacking on acylation rates due to methylation, but there is significant reduction of *k*_acyl_ by C-1 carbon substitution. A similar and somewhat larger effect was seen for the cU modification, which decreased *k*_acyl_ by over 6 fold as a result of replacing electronegative O with C in the sugar ring. Interestingly, the A-context RNAs showed somewhat lower *k*_acyl_ values compared to the U context, resulted a by 1.9 to 2.6-fold reduction depending on the modification. This is consistent with the prior observation of lower acylation reactivity of polyA as compared with polyU.^44^

**Table 2.** Rate constants of acylation reactions^a^ at the central nucleotide of RNA oligonucleotides without or with modifications.

| U oligonucleotide | $k_{\text{acyl}} (\text{M}^{-1}\text{min}^{-1})$ | A oligonucleotide | $k_{\text{acyl}} (\text{M}^{-1}\text{min}^{-1})$ |
| --- | --- | --- | --- |
| U | $0.97 \pm 0.02$ | U | $0.37 \pm 0.09$ |
| m5U | $0.87 \pm 0.02$ | m5U | $0.36 \pm 0.09$ |
| $\Psi$ | $0.27 \pm 0.07$ | $\Psi$ | $0.092 \pm 0.020$ |
| m1Ψ | $0.27 \pm 0.05$ | m1Ψ | $0.17 \pm 0.04$ |
| cU | $0.15 \pm 0.03$ | cU | $0.055 \pm 0.016$ |

<sup>a</sup>Rate constants of RNA 2'-OH acylation using NAI in pH 7.4 buffer at 37 °C were determined from relative peak intensities measured by mass spectrometry. Standard deviation (SD) values are given in parentheses. See full sequences in Table 1.

Overall, although the correlation is not quantitative, each pairwise comparison of N/O substitution by carbon shows the same qualitative effect on acylation rate as it does in strand cleavage rate (Figure 5B). For example, in the polyU context, acylation is decreased for Ψ as compared to U, and strand cleavage rate is also decreased. The same is true for Ψ versus U in the polyA context, and indeed for the pairwise comparisons of U to cU and m5U to m1Ψ. In all cases, substitution of nitrogen or oxygen by carbon decreases acylation rate and decreases strand scission rate as well.

### Correlations of nucleophilicity and p*K*_a_ of alcohols

Taken together, the above data show clearly that carbon replacement of electronegative atoms near the 2*’*-OH group strongly reduces nucleophilicity as measured both by acylation (with an external electrophile) and strand self-cleavage (an internal electrophile). This is consistent with the notion that the replacement of the electronegative atoms reduces the acidity of the 2*’*-OH group via inductive effects. It is difficult to measure the p*K*_a_ of this group directly, as it is already high (*ca*. 12.5-13.5) in native RNA,^22^ and titrations at high pH are likely to cause rapid strand cleavage. Thus we sought to connect nucleophilicity of the 2*’*-OH with its p*K*_a_ by studying a series of small alcohols with p*K*_a_ values between 12-15 (Figures 5C,D), spanning the range of reported p*K*_a_s of the 2*’*-OH of ribonucleotides.^22,23^ The p*K*_a_ of ethanol is 15.9 at 25 °C, and substitution by electronegative atoms at the 2-carbon is well documented to inductively reduce the p*K*_a_ of the OH group.^45^

We measured acylation kinetics for the alcohols in pH 7.4 buffer at 37 °C, and initial reaction rates (Table 3) were determined by ^1^H-NMR (Figures 5C and S8). The data confirmed an inverse relationship of p*K*_a_ with nucleophilicity as measured by acylation rate; a linear plot of log(*k*_acyl_) against p*K*_a_ showed a slope of −0.94 (Figure S9), documenting an average 9-fold increase of kinetic rate per unit of p*K*_a_ for the alcohols. Thus, the data confirm that these alcohols react with NAI at neutral pH at rates governed by their p*K*_a_ values, and the less acidic they are, the lower their nucleophilicity as measured by acylation rate. The data are entirely consistent with a mechanism of reaction involving the anion form of the alcohol reacting with the electrophile, which we hypothesize is the case for RNA as well.

**Table 3.** Rate constants of acylation reaction for alcohols^a^.

| Alcohol | $k_{acyl}$ (NAI)<br>(M<sup>-1</sup>min<sup>-1</sup>) | $k_{acyl}$ (meNAI)<br>(M<sup>-1</sup>min<sup>-1</sup>) |
| --- | --- | --- |
| 2,2,2-Trifluoroethanol | 1.70 ± 0.61 | 1.50 ± 0.25 |
| 2,2,2-Trichloroethanol | 0.79 ± 0.15 | 0.24 ± 0.18 |
| 2,2-Dichloroethanol | 0.36 ± 0.12 | 0.12 ± 0.03 |
| 2-Chloroethanol | 0.013 ± 0.006 | 0.0049 ± 0.0015 |

<sup>a</sup>Rate constants of acylation in presence of NAI (37 °C) and meNAI (25 °C) in buffer (pH 7.4) solution were derived from <sup>1</sup>H NMR analysis of initial rates.

Previously determined p*K*_a_^22^ and corresponding *k*_acyl_ values^21^ for mononucleotides were fit well in our plot of the smaller alcohols (log(*k*_acyl_) *vs* p*K*_a_, Figures S10, S11, and 5E) using a closely analogous reagent (meNAI) and temperature (25 °C), suggesting that the reactivities of ribonucleotides toward NAI are also well explained by this mechanism. If we assume that p*K*_a_ shifts due to carbon substitution for electronegative atoms were entirely responsible for the relative acylation rates seen for the oligonucleotides (Figures 5E and S8), then the 3.6-fold reduction of 2*’*-OH acylation rate from U to Ψ modification suggests that the carbon substitution causes a *ca*. 0.6 p*K*_a_ unit increase for the 2*’*-OH of Ψ relative to U. The larger 6-fold reduction in acylation rates caused by cU substitution relative to U suggests a somewhat greater increase in p*K*_a_ of approximately 0.9, consistent with the greater electronegativity of oxygen relative to nitrogen.

After this work was completed, a preprint describing relevant studies of dinucleotides containing Ψ and m1Ψ was posted.^46^ Comparing U to Ψ, the authors observed a 2-fold decrease in uncatalyzed cleavage rate and a 10-fold decrease in enzymatic cleavage rate by RNase A, consistent with the current findings albeit considerably smaller in magnitude, possibly due to the small size of the substrates in that study. Molecular dynamics simulations suggested a 2.9 kcal/mol free energy difference for proton transfer from 2*’*-OH for U vs. Ψ, attributed to p*K*_a_ difference. That finding is also generally consistent with the current results, although a 2.9 kcal/mol difference suggests a p*K*_a_ difference of *ca*. 2.1 p*K* units, which is significantly larger than our experimental findings and greater than the differences in documented p*K*_a_s of relevant small alcohols such as dichloroethanol and trichloroethanol. Unlike the current study, the authors observed no methylation-associated difference in enzymatic cleavage rates for m1Ψ versus Ψ, whereas we observe substantial slowing as a result of methylation in two nucleotide contexts.

It should be noted that our data show that acylation rates in the RNA oligomers are similar although not identical to those of the current small alcohols as well as for mononucleotides measured previously; however, comparisons of trends within the context of the polymer (such as U versus Ψ) remain consistent. For example, trichloroethanol (p*K*_a_ 12.8), having three electronegative atoms, is more reactive to acylation than dichloroethanol (p*K*_a_ 13.5), and the 2*’*-OH group of U in RNA (also having three electronegative atoms) is more reactive than that of Ψ (with two electronegative atoms) by a similar factor. Work is ongoing to understand differences between the reactivity of RNA relative to simpler alcohol species of similar p*K*_a_.

## CONCLUSIONS

Taken as a whole, our data show that carbon substitution in ribonucleotides, particularly for the atoms attached at C1*’* of ribose, results in strongly reduced nucleophilicity of the 2*’*-OH group in RNAs, causing slower rates of attack on external electrophiles such as acylating agents, and reduced rates of attack on the neighboring phosphodiester group as well. This lower nucleophilicity in carbon-substituted analogs results in markedly slower RNA strand cleavage rates at the phosphodiester bond 3*’* to the nucleotide where substitution occurs. This has relevance both to native RNAs and to therapeutic RNAs containing Ψ and m1Ψ modifications, providing quantitative data for this protective effect as well as basic understanding of its physicochemical origins. The biological roles of Ψ in cellular RNAs are complex^47^ and are under active study;^48–50^ however, the current data provide clear and quantitative documentation of the protective effect of this modification against both uncatalyzed and enzyme catalyzed cleavage. The data also suggest that broader carbon substitution for O4*’*, N1 and other atoms may be worthy of consideration in the design of future modification of RNAs for therapeutic use. In addition, it seems likely that these modifications could also be combined with other known modifications to favorable effect. Studies have demonstrated that hybrid architectures incorporating both ribose and base modifications have been developed to simultaneously enhance RNA stability and tailor biochemical properties for optimized therapeutic performance.^51,52^

We have further shown that alcohols with similar p*K*_a_ as ribonucleotides have similar nucleophilic reactivity, and that alcohols with reduced number of electronegative atoms have reduced reactivity. In RNA, we show that this also the case with Ψ and m1Ψ relative to their N-nucleoside counterparts U and m5U. Thus, although direct p*K*_a_ measurements of 2*’*-OH in RNA are not yet available, the data suggest that the C-nucleosides Ψ and m1Ψ stabilize RNA by reducing the acidity of their associated 2*’*-OH groups as compared with canonical uridine, due to reduced inductive stabilization of the anionic form of the 2*’*-OH group.

In addition to effects on 2*’*-OH acidity and nucleophilicity, our data also show that methyl substitution of bases at C-5 of U and analogously at N1 of Ψ can also have protective effects on stabilizing phosphodiester bonds in RNA, depending on the context. Both of these substitutions enhance base stacking, which seems likely to be relevant to explaining their protective effect, as the substitution is too remote from the 2*’*-OH to cause a direct effect on reactivity. We observe that the effect for spontaneous cleavage is mixed (depending on context, as is the case for stacking generally^20^) and requires more study before its origins and scope can be fully understood. However, the effect has not been reported previously and is worthy of consideration in the biological context. The analogous C-5 methylation of cytosine in RNA is known^40^ and the current results suggest that, like m5U, it has the potential to affect the stability of adjacent phosphodiester bonds. Interestingly, the adenine modification m6A, also widely observed in cellular RNA, is also known to enhance base stacking,^53^ and recent experiments have shown that it does indeed affect the reactivity of adjacent nucleotides substantially, enabling the use of acylating agents to detect positions of m6A in cellular RNA.^54^ Thus, the current data suggest that these other methyl substitutions in native RNAs bear further study in light of possible alterations of nearby 2*’*-OH reactivity.

## ASSOCIATED CONTENT

### Supporting Information

The supporting information is available at https://, and includes experimental methods and materials, gel images, plots of UV-melting and RNA decay data, and CD, MALDI-TOF, and ^1^H-NMR spectra.

## ACKNOWLEDGEMENTS

We thank the U.S. National Institutes of Health (GM145357) for support.

