## supplementary materials for "C-Nucleosides Stabilize RNA by Reducing Nucleophilicity at 2’-OH" (from another version of this preprint: version 1, 2025-07-27)

### Materials and Methods

#### Materials

eGFP mRNAs were purchased from TriLink Biotechnology. Short RNA oligonucleotides were purchased from IDT or synthesized by Stanford PAN facility unless otherwise stated (Supplementary Table S1). Purchased RNAs were diluted with nuclease-free water and concentrations measured with a Thermo Scientific Nanodrop spectrophotometer. All chemicals were purchased from commercial suppliers and used without further purification (Supplementary Table S2). All enzymes, kits, bio-reagents and software were obtained from the sources listed in Supplementary Table S2.

#### Accelerated mRNA ageing experiments

Two mRNAs (eGFP and m1ΨeGFP) were used to evaluate the effect of base modification on mRNA thermal stability. 20 $\mu$L of a solution containing 10 ng $\mu L^{-1}$ of mRNA species in RNase-free water was incubated in a sealed RNase-free PCR tube at 37 °C. After the desired time, 3 $\mu$L of the mRNA solution was collected at each time point and RNA integrity was analysed by Capillary Electrophoresis (CE) on Agilent RNA 6000 Pico Chips using an Agilent 2100 Bioanalyzer. CE data were processed and visualized using 2100 Expert software (Agilent). CE traces are shown in Figure 2a. The apparent half-lives for thermal degradation were determined from the slope values of the linear relations by plotting the natural logarithm of the parent peak intensity from CE traces against time.

**UV-melting experiments**

UV absorbance was measured by using a Varian Cary 300 UV-visible spectrophotometer equipped with thermoprogrammer. The melting curves (absorbance versus temperature) of the oligonucleotides at 5 $\mu$M in 1x PBS buffer solutions without or with 10 mM $MgCl_2$ were obtained at 260 nm by annealing from 70 to 10 °C, followed by heating from 10 to 70 °C at a rate of 1 °C $min^{-1}$.

**PAGE analysis of RNA thermal degradation**

For kinetics assays of thermal degradation, RNA oligonucleotides (5-120 $\mu$M) were incubated for varied times in a reaction buffer containing a final concentration of 50 mM CHES buffer (pH 10.0) and 10 mM $Mg^{2+}$.[S1] Temperatures were maintained at 37 °C unless otherwise indicated. Reaction aliquots (10 $\mu$L) at each time point were quenched by diluting into 2.5 $\mu$L 1 M Tris-HCl (pH 7.0) and 1 $\mu$L 0.5 M EDTA. Quenched samples were further diluted (2-fold) by adding to 8 M urea loading buffer with dye and stored at –80 °C prior to gel loading. Reaction mixtures were separated by 20% denaturing PAGE in 1x TBE buffer (pH 8.3) at 25 mA for 1 h and the gel image was acquired by iBright 1500 imager after staining with SYBR-gold. After converting images to 16-bit using ImageJ software, analysis was done by plotting band intensities from same sized rectangular area on selected bands. Quantified band intensities were used to calculate remaining RNA fractions for measuring kinetic parameters.

### RNA enzymatic degradations

For kinetics assays of enzymatic degradation, RNA oligonucleotides (100 $\mu$M) were incubated at 37 °C for 10 min in 30 $\mu$L reaction buffer containing 100 mM Tris-HCl buffer (pH 7.4). 30 $\mu$L of 20 ng $mL^{-1}$ RNase A solution was then added and the mixture incubated for varied reaction time. Reaction aliquot (10 $\mu$L) at each time point was quenched by diluting into 1 $\mu$L RNaseOUT$^{TM}$ (40 U$\mu L^{-1}$), 1 $\mu$L of 0.5 M EDTA, and 1 $\mu$L of 100 mM DTT. Quenched samples were further diluted (2-fold) by adding to 8 M urea loading buffer with dye and storing at –80 °C prior to gel loading. Reaction mixtures were separated by 20% denaturing PAGE and analyzed by using ImageJ software following the method described above. The rate constants of pseudo-first-order ($k_{clv(app)}$) and second-order ($k_2$) kinetics were quantified (Table S3) following the method described below.

### Determination of rate constants

Rate constants were determined under various reaction conditions by first plotting the natural logarithm of the fraction of substrate remaining uncleaved at different incubation times. The negative slope of the resulting line was generated by a least-squares fit to the data, reflecting the rate constant for RNA cleavage by transesterification. Data reported are averages of at least three separate experiments.

### CD measurements

Circular Dichroism (CD) spectra were obtained using a JASCO-820 spectropolarimeter with a temperature controller. The CD spectra were collected from 350 to 200 nm wavelength at a scan rate of 50 nm min$^{-1}$ in 0.1 cm path-length cuvette at 4 °C. All spectra were measured at a strand concentration of 20 $\mu$M in 1x PBS buffer (pH 7.4) solution with 10 mM $MgCl_2$. The cuvette-holding chamber was continuously flushed with a stream of dry $N_2$ gas to avoid water condensation on the cuvette exterior.

**Synthesis of acylimidazole reagents**

Reagents (NAI and meNAI) were synthesized following the general protocol.[S2,3] The carboxylic acid precursor (1.0 equiv.) was dissolved in dry DMSO as a 4 M solution. To it was added an equal volume of suspension containing 1.0 equiv. 1,1′-carbonyldiimidazole in dry DMSO at room temperature. The resulting solution was stirred at room temperature for 1 (NAI) or 3-4 h (meNAI). The completion of reaction was monitored by $^1$H NMR spectroscopy, which confirmed a 1:1 mixture of the acylimidazole and free imidazole. After the reaction, the crude solutions were used as a 2 M acylimidazole stock solution without further purification.

**Kinetics studies of RNA acylation**

1.0 $\mu$L of RNA stock (100 $\mu$M) was dissolved in 8.5 $\mu$L water solution with 1x PBS buffer (pH 7.4) and incubated at 37 °C for 15 min. Then, 0.5 $\mu$L freshly prepared 2 M acylating reagent was added to RNA solution to kept DMSO 5%. The reaction was incubated at 37 °C for the indicated time and quenched by diluting with RNase-free water to 500 $\mu$L. The mixture of RNA and acylated RNA was purified with Amicon ultra centrifugal filtration (3K) and stored at –80 °C. The ratio of substrate and product in the RNA mixture (as peak intensities) was analyzed with MALDI-TOF operating in the linear negative mode in 3-HPA matrix.

### Mass spectrometry

The MALDI-TOF mass spectra were recorded on a Bruker Daltonik Microflex MALDI-TOF spectrometer with a $N_2$ laser, in linear negative mode. An MSP BigAnchor 96 ground steel target was used for MALDI-TOF analysis. For the linear negative mode, a mixture of 0.3 M trihydroxyacetophenone in EtOH and 0.1 M aqueous ammonium citrate (2:1) used as matrix/comatrix, and always freshly prepared before analysis. Spectral data were analyzed with Flex Analysis, version 3.4 (Bruker).

### The p$K_a$ determination of alcohols

Four alcohols (2,2,2-trifluoroethanol, 2,2,2-trichloroethanol, 2,2-dichloroethanol, 2-chloroethanol were selected to study acylation reaction. The p$K_a$ values of selected alcohols were reported as 12.4, 12.2, 13.1, and 14.3, respectively.[S4] Reported p$K_a$ values were used in this study.

### Kinetics study for the acylation of alcohols

1 M alcohol solutions were prepared with selected alcohols in $D_2O$. For 200 $\mu$L reaction mixture, 20-40 $\mu$L 1 M alcohol solution was mixed with $D_2O$ and 10 $\mu$L solution of 20x PBS buffer (pD 7.4) prepared in $D_2O$. The mixture was incubated at 37 °C or 25 °C for 15 min. Then, to the mixture 10-20 $\mu$L 2 M stock solution of the acylating reagents in DMSO-$D_6$ was

added to react with alcohols in 1:1 ratio and kept DMSO 5-10%. The reaction was incubated at 37 °C or 25 °C for the indicated time and quenched by diluting with RNase-free water to 700 $\mu$L. The $^{1}$H-NMR of reaction mixture was immediately recorded after quenching the reaction and analyzed the reaction rate from the ratio of substrate and product in the reaction mixture (as peak intensities).

**NMR spectroscopy**

$^{1}$H NMR spectra were recorded on Varian Mercury 500 MHz NMR spectrometer. Spectra were internally referenced to the residual solvent signal.

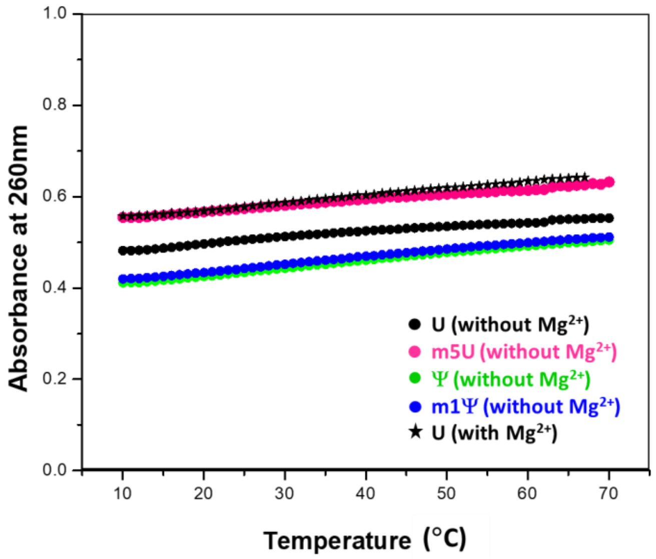

**Figure S1.** UV thermal denaturation plots at 5 $\mu$M concentration for A oligonucleotides containing U (black), m5U (pink), Ψ (green), and m1Ψ (blue) in 1x PBS buffer solutions without (circle) and with (star) 10 mM $MgCl_2$.

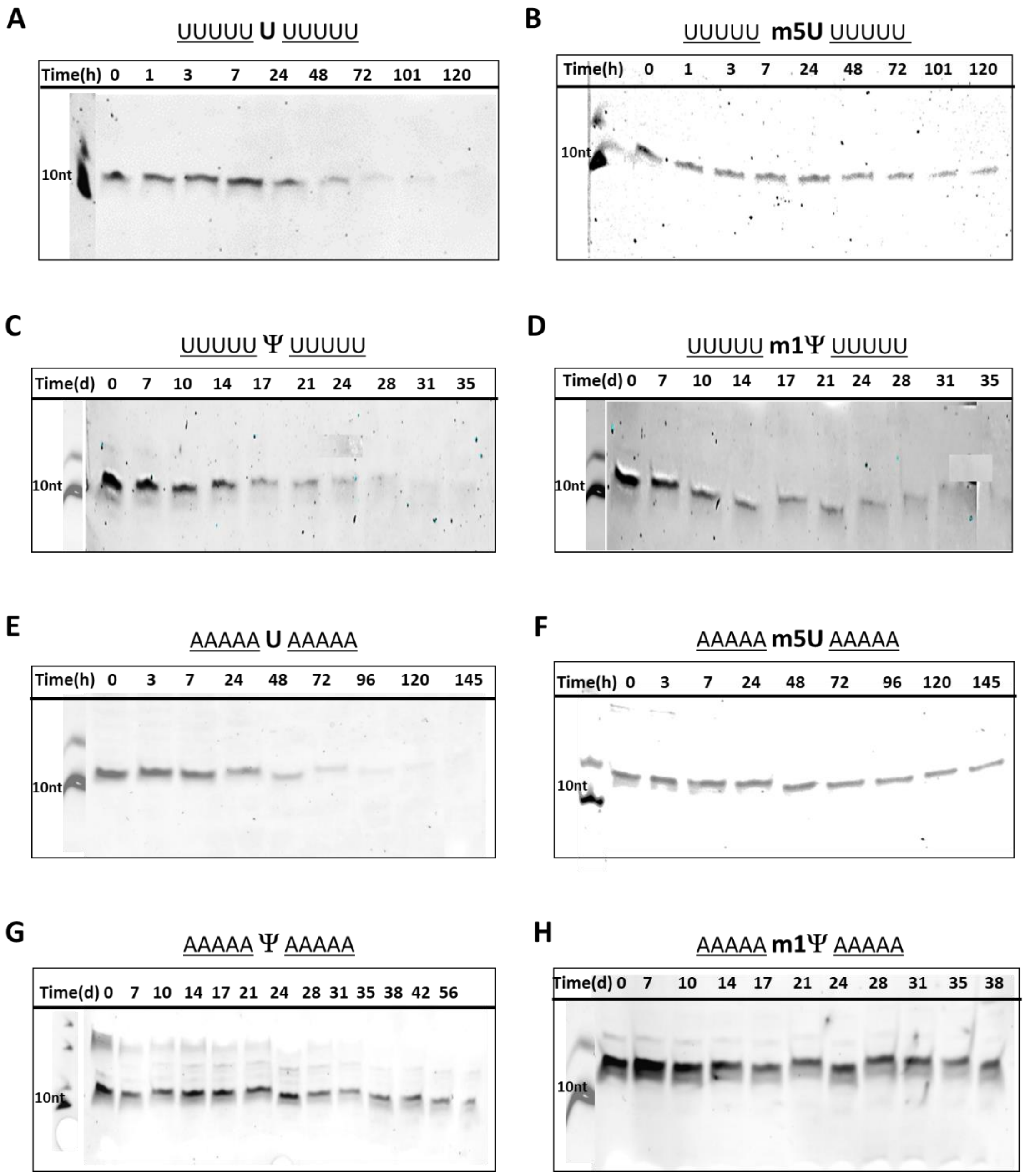

**Figure S2.** Representative gel images (20% denaturing PAGE) for thermal degradation of U, m5U, Ψ, and m1Ψ in U contexts (A, B, C, D respectively) U and A contexts (E-H, respectively) after incubating substrate oligonucleotides in CHES buffer (pH 10.0) solution with $Mg^{2+}$ at 37 °C for the noted time period.

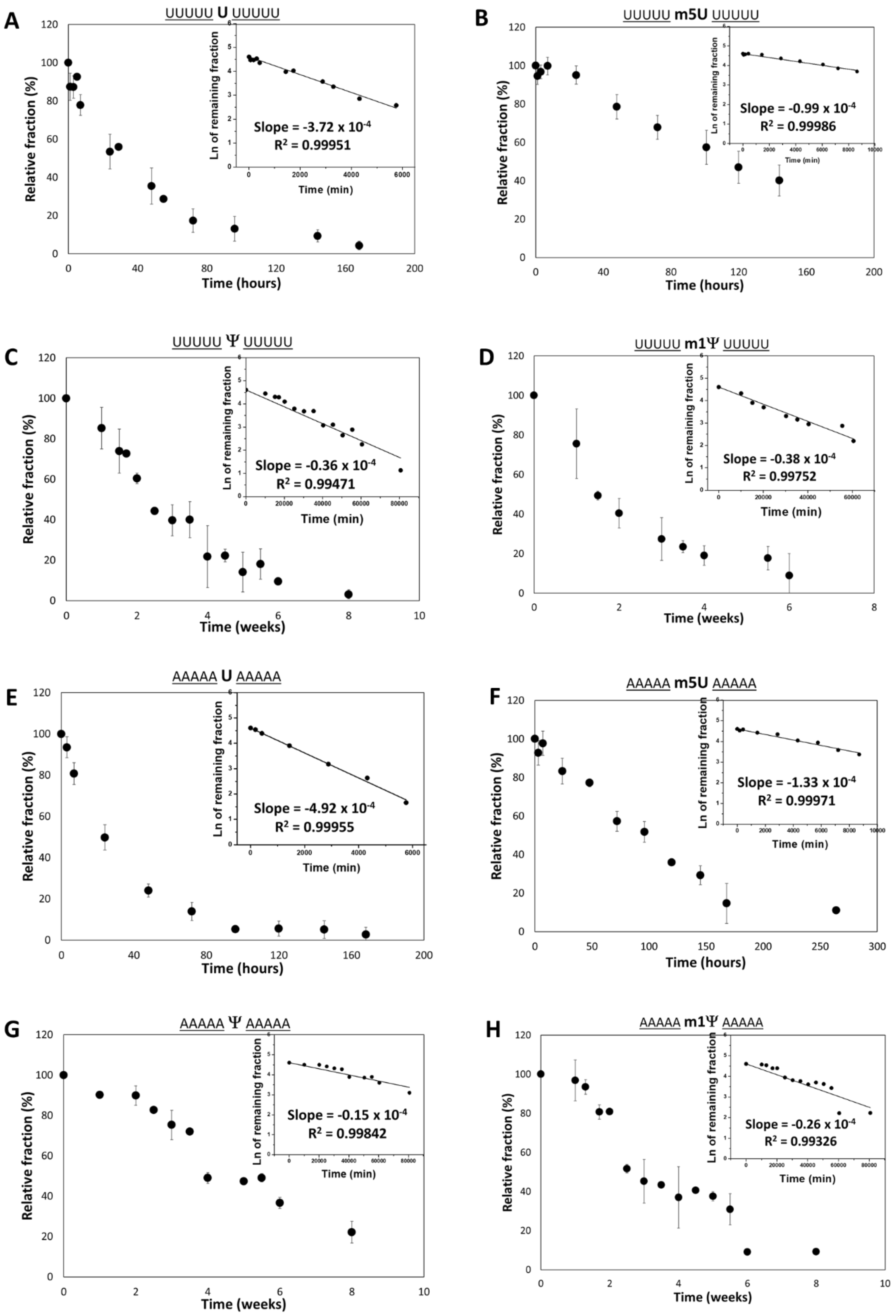

**Figure S3.** Plots of the decay of the central RNA linkage for U, m5U, Ψ, and m1Ψ in U contexts (A, B, C, and D, respectively) and A contexts (E, F, G, and H, respectively). The relative intensity of bands was derived from gel analysis (see examples in Fig. S2). Negative slope of each linear plot reflects the rate constant $k_{clv}$ ($min^{-1}$) for thermal degradation of the oligonucleotide in the CHES buffer (pH 10.0) at 37 °C.

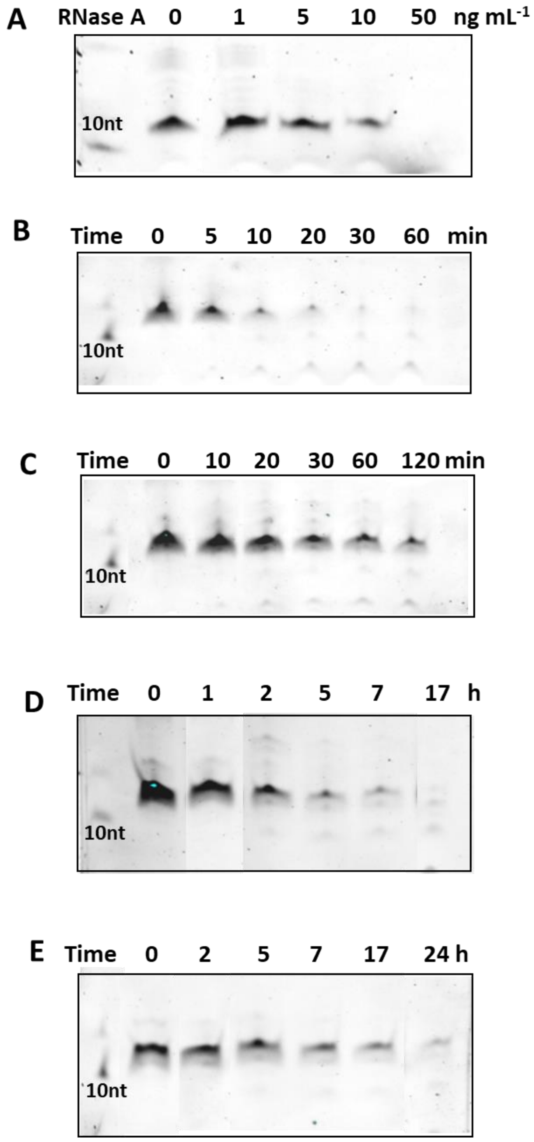

**Figure S4.** A. Representative denaturing 20% PAGE image analyzing ribonuclease A catalyzed cleavage for screening of RNase A concentrations as shown, with 50 µM unmodified A context oligonucleotide using RNase A at 0 to 50 ng $mL^{-1}$ concentration in final reaction mixture after 10 min reaction. B, C, D, E. Representative images of 20% denaturing PAGE gels for for thermal degradation of U, m5U, Ψ, and m1Ψ, respectively, in A contexts after incubating substrate oligonucleotides with 10 ng $mL^{-1}$ in Tris-HCl buffer (pH 7.4) solution at 37 °C for the noted time.

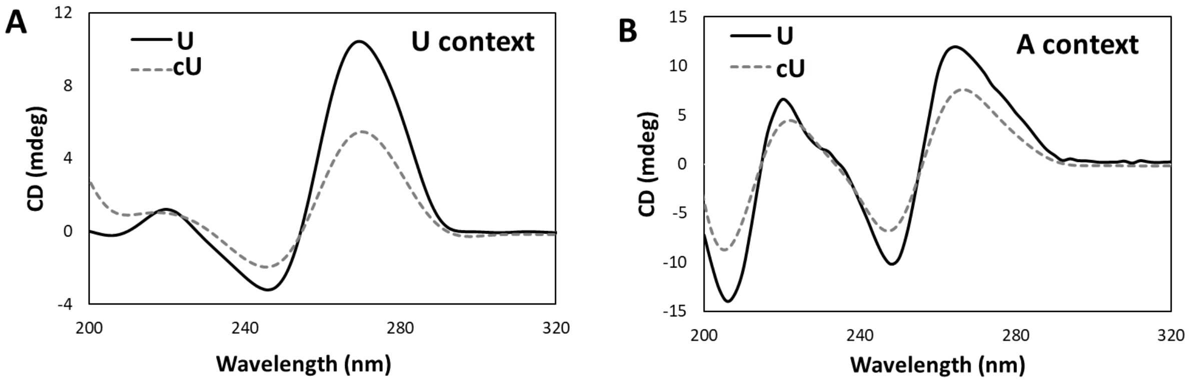

**Figure S5.** Comparison of CD spectra of U and A oligonucleotides, respectively, containing cU modification with corresponding unmodified oligonucleotides of 20 µM concentration at 4 °C in the 1xPBS buffer (pH 7.4) containing 10 mM $Mg^{2+}$.

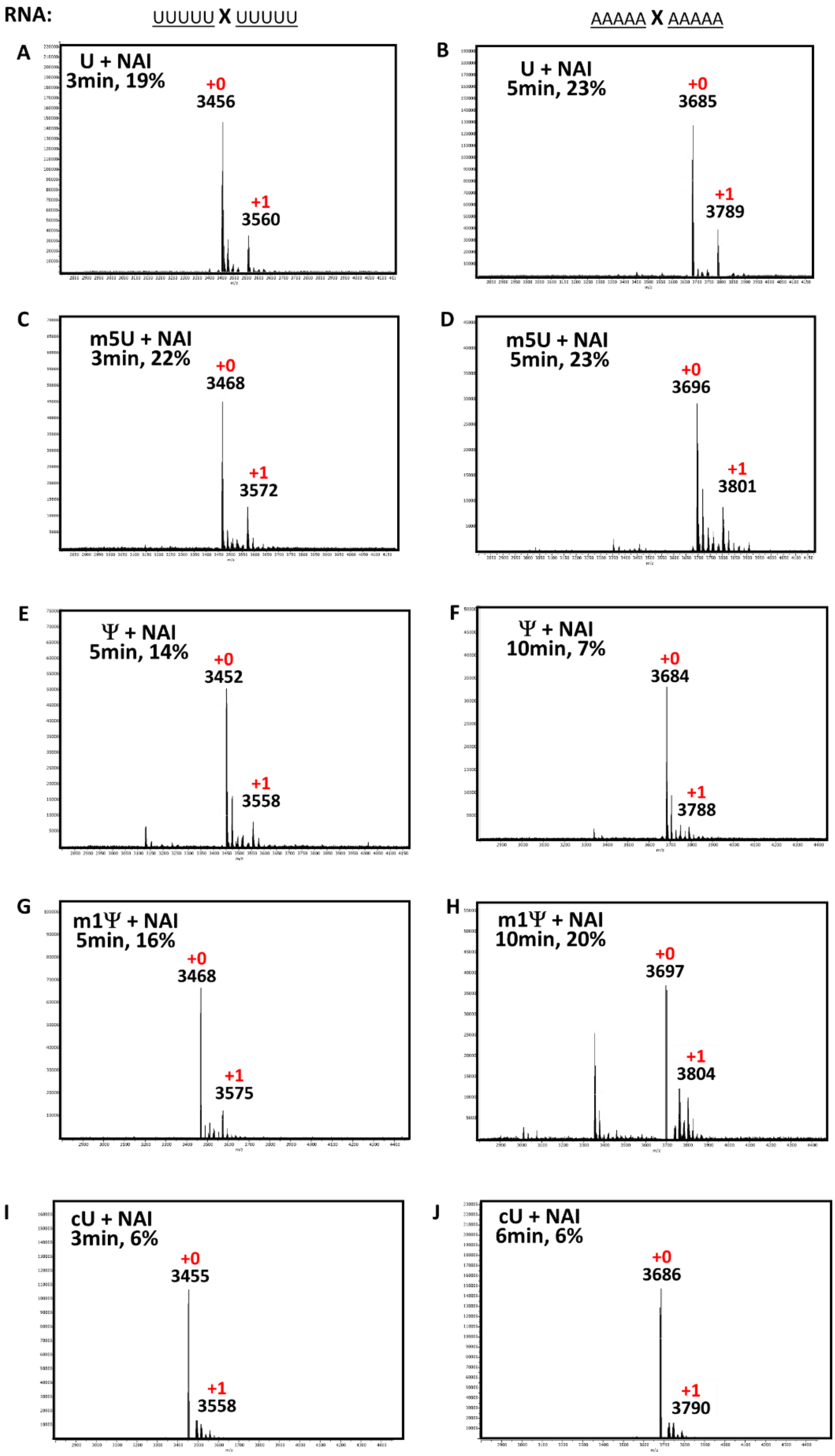

**Figure S6.** Representative MALDI-TOF spectra after indicated times of acylation reactions with NAI (100 mM) for RNA oligomers (10 $\mu$M) with uridine modifications (X), such as U, m5U, Ψ, m1Ψ, and cU in (A, C, E, G, and I, respectively) U and (B, D, F. H, and J, respectively) A contexts at 37 °C to evaluate kinetics of 2'-OH reactivity shown in Table 2.

### $^{1}H$ NMR spectrum of TFE after 2min reaction (25% conversion)

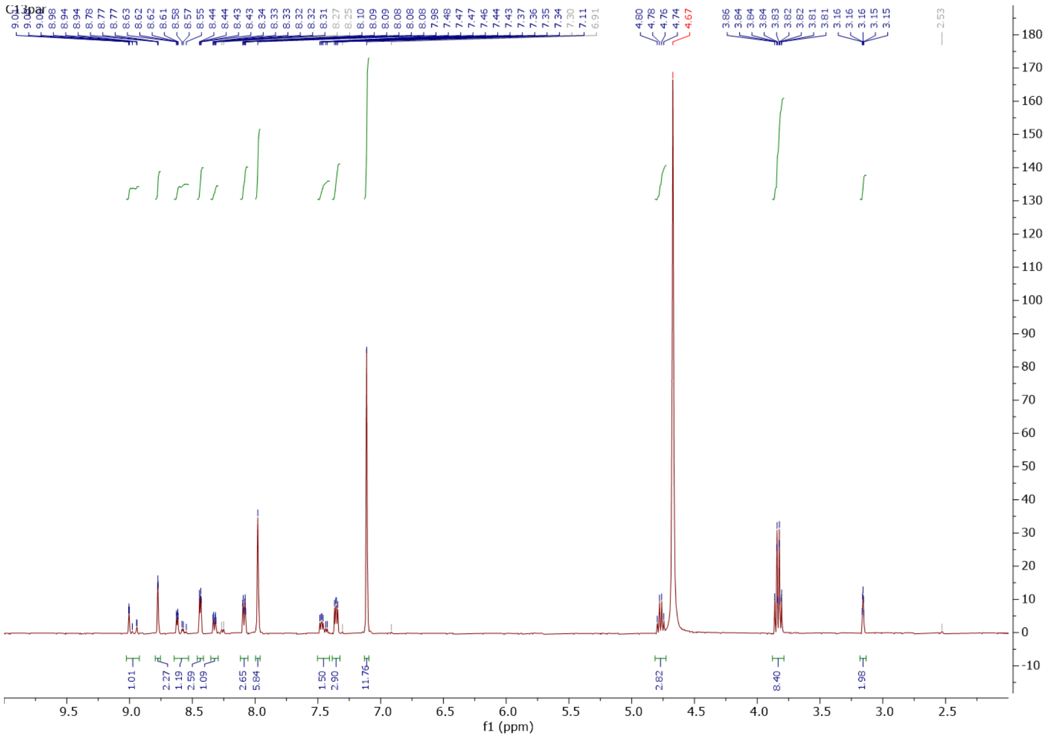

### $^{1}H$ NMR spectrum of TCE after 3min reaction (13% conversion)

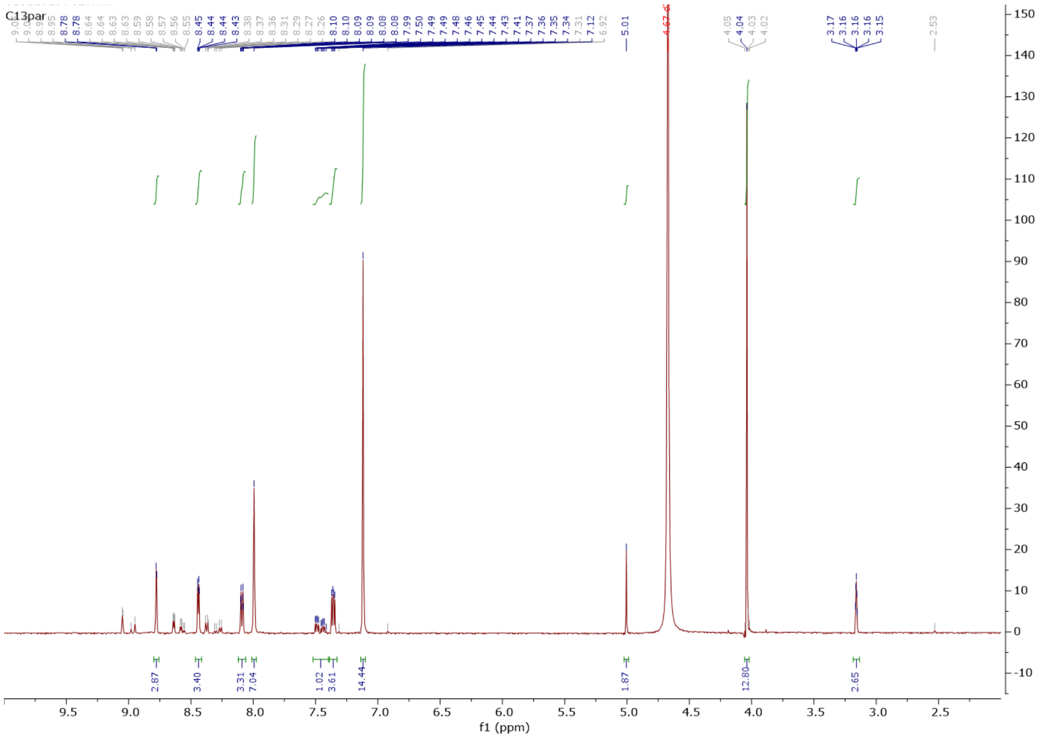

(Figure S7 continued)

$^1H$ NMR spectrum of DCE after 8min reaction (18% conversion)

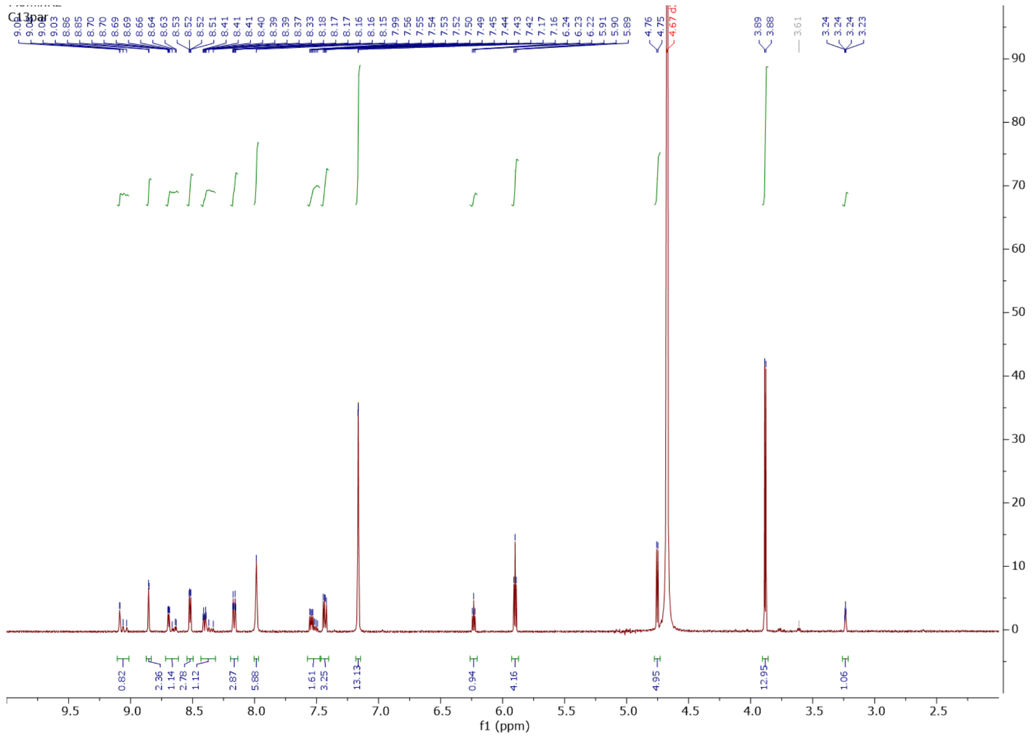

$^1H$ NMR spectrum of CE after 20min reaction (10% conversion)

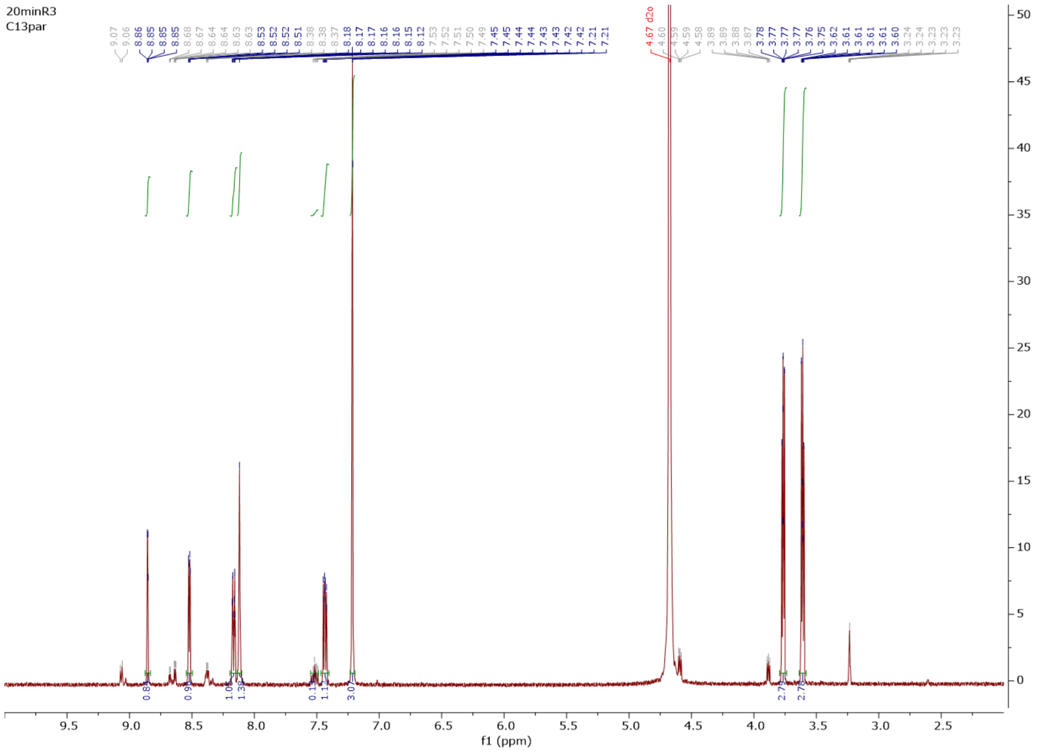

**Figure S7.** Representative $^1H$ NMR spectra after indicated time of acylation reaction with NAI and alcohols (1:1) at 37 °C to evaluate kinetics of 2′-OH acylation. 2,2,2-trifluoroethanol, 2,2,2-trichloroethanol, 2,2-dichloroethanol, and 2-chloroethanol alcohols are written as TFE, TCE, DCE, and CE, respectively. The intensities of corresponding peaks of substrate and products were used to analyze rate constants of second-order kinetics in Table 3 and Figure S8.

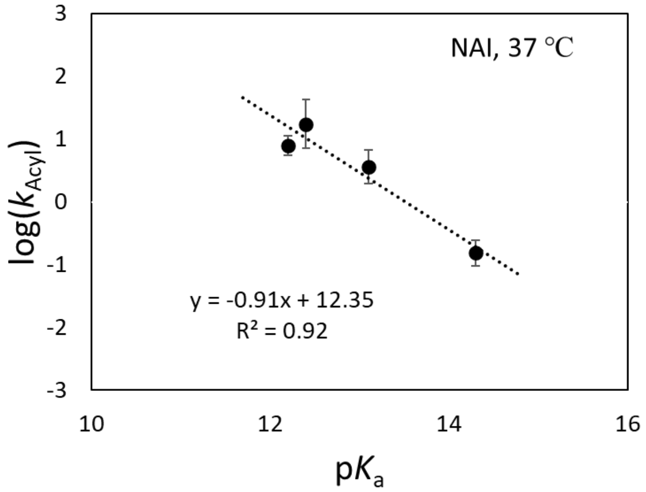

**Figure S8.** Plot of log values for the acylation rate constants ($k_{acyl}$) with NAI versus p$K_a$ for primary alcohols (see Figure 5E) (black circles) at 37 °C for evaluating the contribution of p$K_a$ shift to the reactivity of hydroxyl groups.

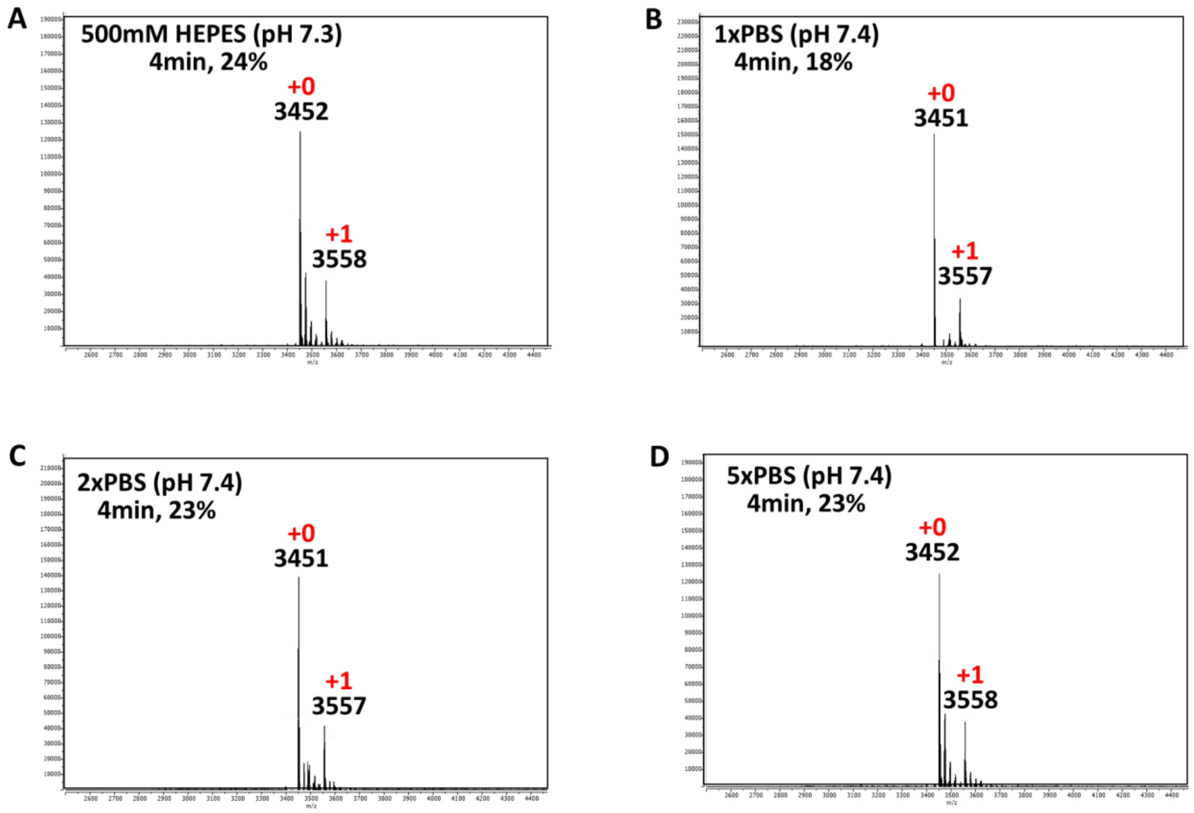

**Figure S9.** Representative MALDI-TOF spectra of after 4 min of acylation reaction with NAI (100 mM) for unmodified U context (10 $\mu$M) at 37 °C in the solution of A. 500 mM HEPES (pH 7.3), B. 1x, C. 2x, and D. 5x PBS buffer (pH 7.4), showing independency of 2'-OH acylation reaction on the buffers.

### $^1H$ NMR spectrum of TFE after 4min reaction (40% conversion)

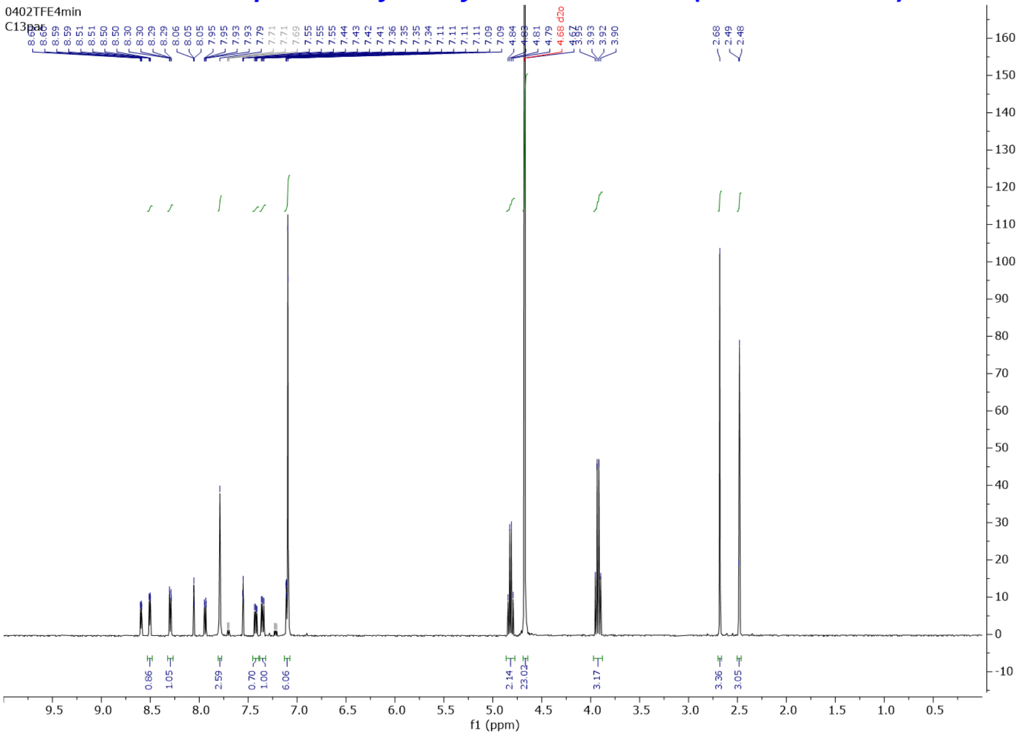

### $^1H$ NMR spectrum of TCE after 6min reaction (31% conversion)

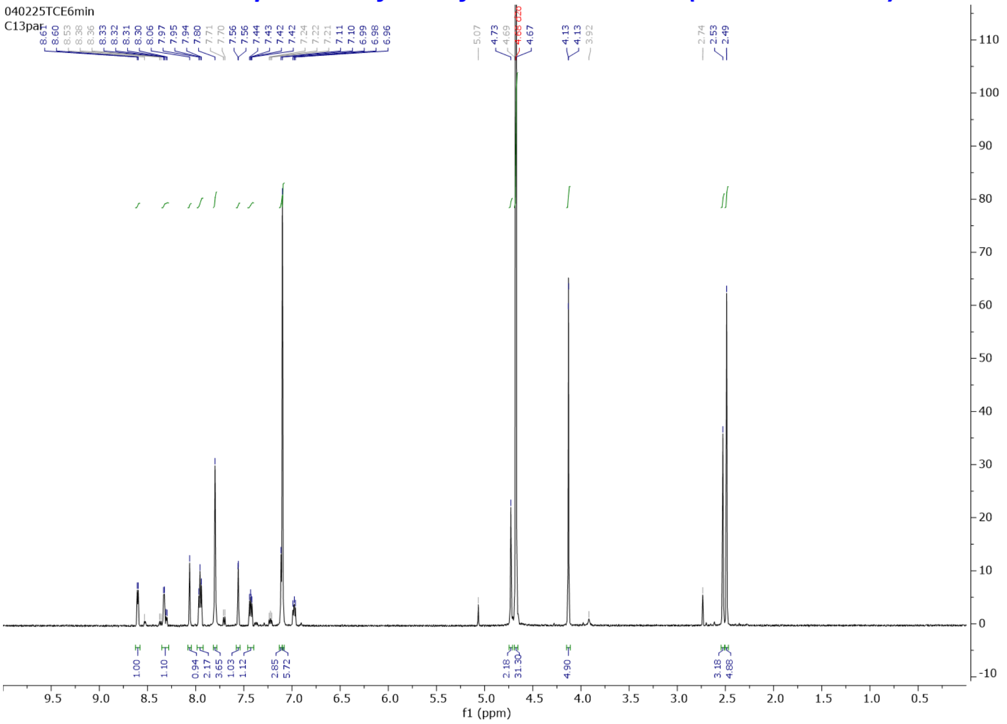

(Figure S10 continued)

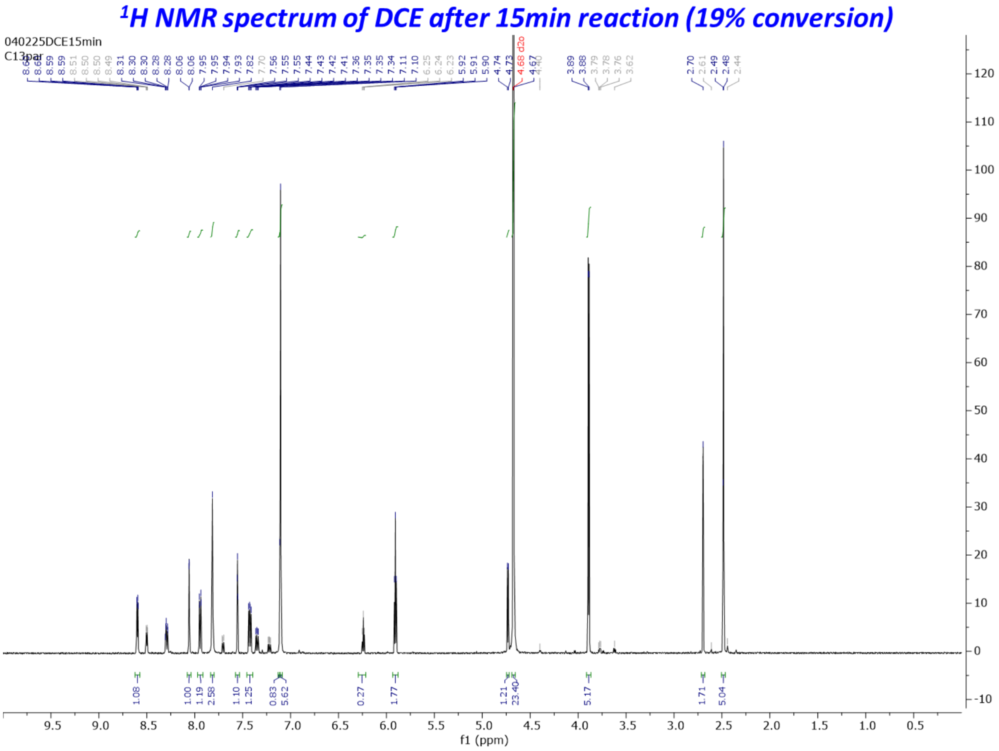

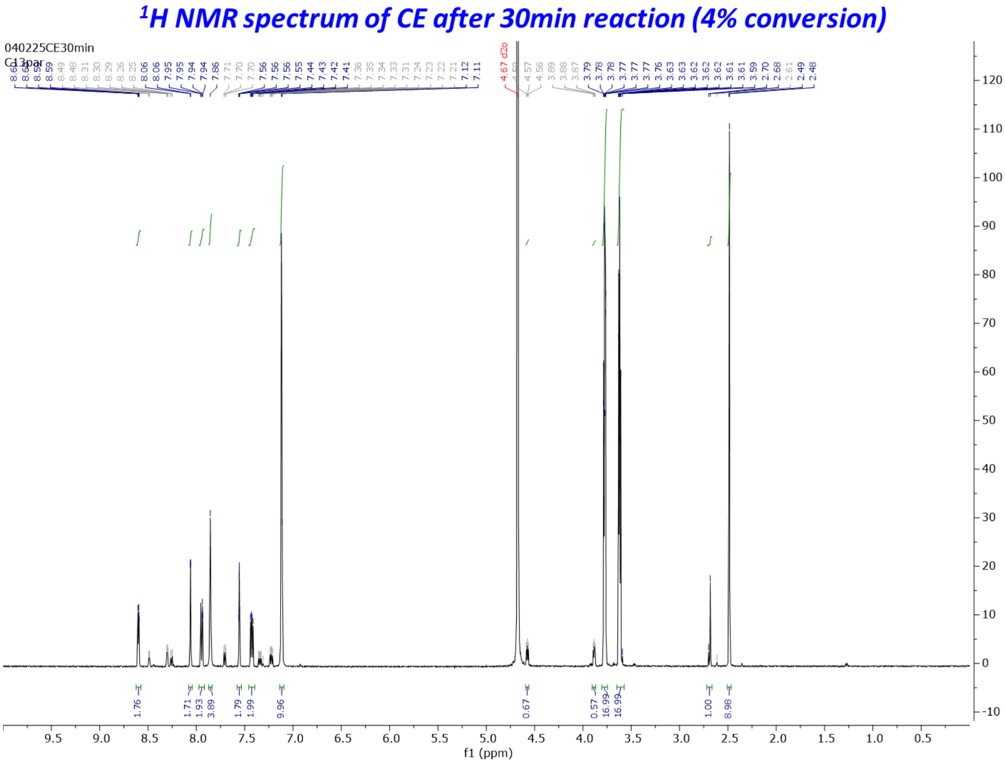

**Figure S10.** Representative $^1$H NMR spectra after indicated times of acylation reaction with meNAI and alcohols (1:1) at 25 °C to evaluate kinetic order of corresponding OH acylation. Here, 2,2,2-trifluoroethanol, 2,2,2-trichloroethanol, 2,2-dichloroethanol, and 2-chloroethanol alcohols are written as TFE, TCE, DCE, and CE, respectively. The intensities of corresponding peaks of substrate and products were used to analyze rate constant of second-order kinetics in Table 3 and Figure 6C.

**Table S1:** List of short oligonucleotides used in this study

| **RNA Sequence[a] (5′→3′)** | **Source** |
|---|---|
| UUUUU **U** UUUUU | IDT (Standard desalting) |
| UUUUU **m5U** UUUUU | Stanford-PAN facility (HPLC purified) |
| UUUUU **Ψ** UUUUU | Stanford-PAN facility (HPLC purified) |
| UUUUU **m1Ψ** UUUUU | Stanford-PAN facility (HPLC purified) |
| UUUUU **cU** UUUUU | Alnylam (HPLC purified) |
| AAAAA **U** AAAAA | IDT (Standard desalting) |
| AAAAA **m5U** AAAAA | Stanford-PAN facility (HPLC purified) |
| AAAAA **Ψ** AAAAA | Stanford-PAN facility (HPLC purified) |
| AAAAA **m1Ψ** AAAAA | Stanford-PAN facility (HPLC purified) |
| AAAAA **cU** AAAAA | Alnylam (HPLC purified) |

[a]RNA of 11nt of written sequences were used containing 2′-O-methyluridine (U), uridine (**U**), 5-methyluridine (**m5U**), pseudouridine (**Ψ**), N1-methylpseudouridine (**m1Ψ**), carbocyclic uridine (**cU**), 2′-O-methyladenosine (A).

**Table S2.** List of reagents and materials

| Reagents | Source | Identifier |
|---|---|---|
| **Reagents, Buffers, and Enzymes** | | |
| 5 M NaCl | Invitrogen$^{TM}$ | #AM9760G |
| 1 M $MgCl_2$ | Invitrogen$^{TM}$ | #AM9530G |
| CHES, pH=10.0 | Boston Bioproduct, Inc. | #BB-87 |
| HEPES, pH=7.3 | Gibco$^{TM}$ | #15630080 |
| 1xPBS, pH=7.4 | Gibco$^{TM}$ | #10010-023 |
| 10xTBE buffer | Invitrogen$^{TM}$ | #15581044 |
| Urea | Fisher bioreagents | #BP169-500 |
| 40wt% Acrylamide-bisacrylamide | Sigma-Aldrich | #A9926-100ML |
| UltraPure DTT | Invitrogen$^{TM}$ | #15508013 |
| N, N, N′, N′-Tetramethyl ethylenediamine | Sigma-Aldrich | #110732 |
| Ammonium Persulfate | Thermo Scientific$^{TM}$ | #17874 |
| SYBR Gold Nucleic acid gel stain (10,000x) | Thermo Scientific$^{TM}$ | #S11949 |
| RNase A | Thermo Scientific$^{TM}$ | #EN0531 |
| RNaseOUT$^{TM}$ Recombinant Ribonuclease Inhibitor | Invitrogen$^{TM}$ | #10777019 |
| UltraPure DNase/RNase-Free Distilled water | Invitrogen$^{TM}$ | #10977015 |
| 96% EtOH | Thermo Scientific$^{TM}$ | #T032021000CS |
| RNA Gel loading Dye (2x) | Thermo Scientific$^{TM}$ | # R0641 |
| 3 M sodium acetate, pH=5.5 | Invitrogen$^{TM}$ | # AM9740 |
| Glycogen, RNA grade | Thermo Scientific$^{TM}$ | #R0551 |
| **Commercial kits** | | |
| Amicon Ultra-0.5 3K-Centrifugal Filter Unit | Millipore-Sigma | # UFC500324 |
| Corning Costar Spin-X centrifuge filters, 0.22 µm | Millipore-Sigma | |
| RNA clean-up and concentrator-5 column | Zymo Research | # R1016 |
| **Chemicals and Solvents** | | |
| 1,1’-Carbonyldiimidazole (CDI) | TCI | #C0119 |
| Nicotinic acid | AK Scientific | #J52008-25g |
| 2-Methylnicotinic acid | TCI | #M2089 |
| DMSO | Fisher bioreagents | #BP231-100 |
| D6-DMSO | CIL, Inc. | #DLM-10-10 |
| D2O | CIL, Inc. | #DLM-4-100 |
| Methanool-D4 | CIL, Inc. | #DLM-24-10 |
| Methanol | Fisher Chemical | #A412-500 |
| 2,2,2-Trifluoroethanol | Thermo Scientific$^{TM}$ | #139751000 |
| 2,2,2-Trichloroethanol | Sigma-Aldrich | #T54801-100G |
| 2,2-Dichloroethanol | Chem Cruz$^{R}$ | #SC-230697 |
| 2-Chloroethanol | Sigma-Aldrich | #185744-50G |
| 1,3-Dichloro-2-ethanol | TCI | #D0402 |
| Diacetin | Sigma-Aldrich | #W500615 |
| Deuterium Chloride 35wt% solution | Sigma-Aldrich | #543047-10G |

| | | |
|---|---|---|
| Sodium deuteroxide 40wt% solution | Sigma-Aldrich | #372072-10G |
| Sodium phosphaste monobasic | Sigma-Aldrich | #S5011-100G |
| Sodium phosphaste dibasic | Sigma-Aldrich | #S9763-100G |
| **Software** | | |
| Typhoon Scanner | GE healthcare | NA |
| ChemDraw | Revvitycloud | NA |
| ImageJ | NIH | NA |
| MestReNova | Mestrelab | NA |

**Table S3.** Rate constants for enzymatic cleavage[a] at the central nucleotide of RNA oligonucleotides with RNase A.

| A oligonucleotide | $k_{clv(app)}$ ($min^{-1}$) | $k_2$ ($M^{-1}min^{-1}$) |
|---|---|---|
| U | 28.0 (1.01) x$10^{-2}$ | 384.0 ± 13.8 |
| m5U | 1.2 (0.07) x$10^{-2}$ | 16.4 ± 0.959 |
| Ψ | 0.53 (0.05) x$10^{-2}$ | 7.26 ± 0.685 |
| m1Ψ | 0.26 (0.01) x$10^{-2}$ | 3.56 ± 0.0822 |

[a]Rate constants derived from 20% PAGE analysis. Conditions: Tris-HCl buffer (50 mM, pH 7.4) at 37 °C. Data are averages of 3 replicates; error as standard deviations are given in parentheses. See full sequences in Table 1.
